# Uncovering context-specific genetic-regulation of gene expression from single-cell RNA-sequencing using latent-factor models

**DOI:** 10.1101/2022.12.22.521678

**Authors:** Benjamin J. Strober, Karl Tayeb, Joshua Popp, Guanghao Qi, M. Grace Gordon, Richard Perez, Chun Jimmie Ye, Alexis Battle

## Abstract

Genetic regulation of gene expression is a complex process, with genetic effects known to vary across cellular contexts such as cell types and environmental conditions. We developed SURGE, a method for unsupervised discovery of context-specific expression quantitative trait loci (eQTLs) from single-cell transcriptomic data. This allows discovery of the contexts or cell types modulating genetic regulation without prior knowledge. Applied to peripheral blood single-cell eQTL data, SURGE contexts capture continuous representations of distinct cell types and groupings of biologically related cell types. We demonstrate the disease-relevance of SURGE context-specific eQTLs using colocalization analysis and stratified LD-score regression.

## Background

A complete, mechanistic understanding of the genetic basis of complex traits could provide insights into the biological basis of human health and disease. A powerful approach to filling in the missing links between genetics and complex traits is to use molecular measurements, such as gene expression levels, as an intermediate phenotype. Genetic variants significantly associated with gene expression are known as expression quantitative trait loci (eQTLs) [1–5]. Although eQTL studies have now been performed in large cohorts and numerous tissues [5,6], characterizing the impact of regulatory genetic variants is far from complete. This complexity arises in part because the effects of genetic variation on gene expression vary considerably between different cellular contexts, such as cell types, developmental stage, or condition (Fig. 1A) [7–14].

**Figure 1:**
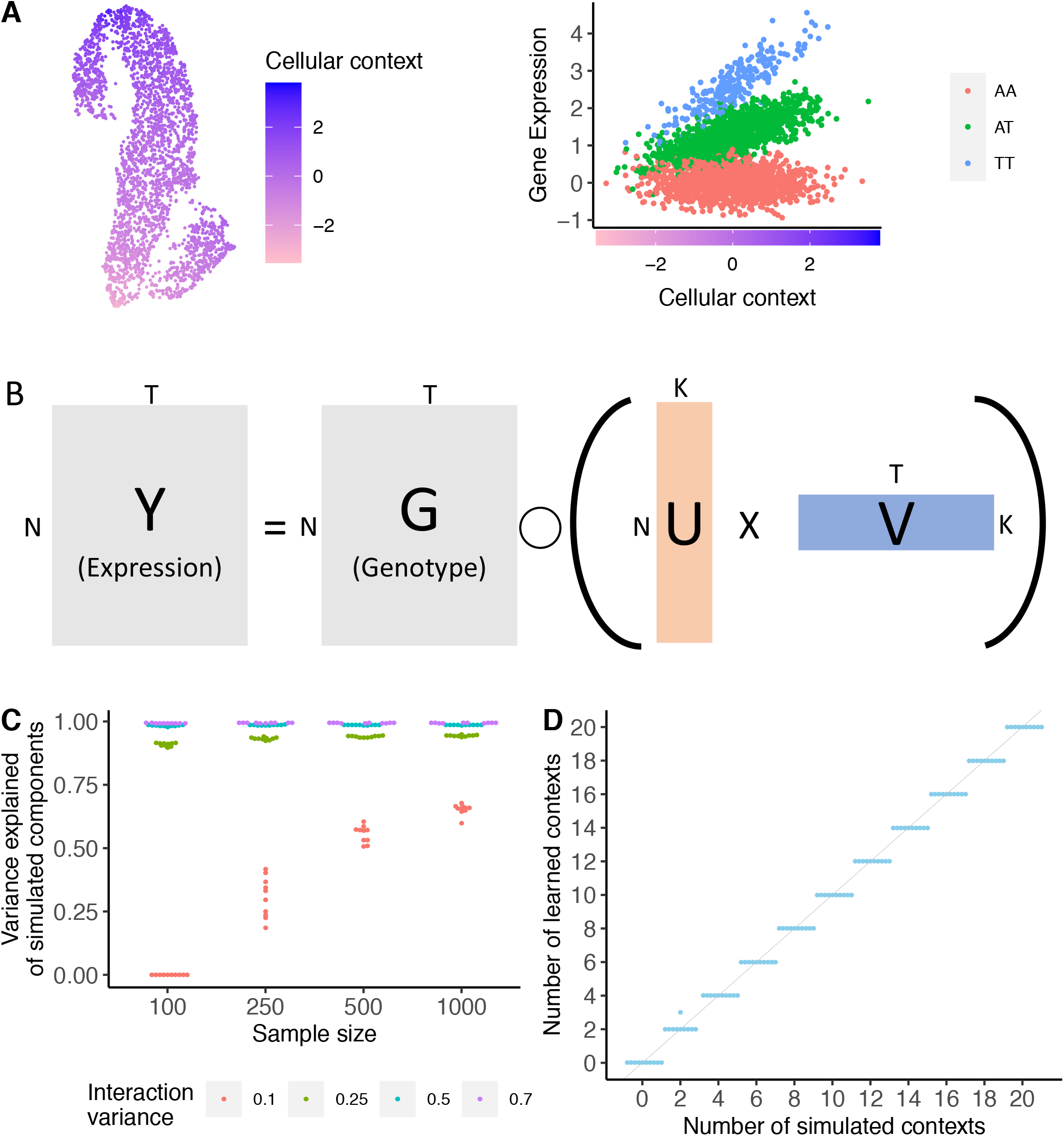
SURGE model overview and simulation: (A) Schematic example of an interaction eQTL where the eQTL effect size (right) changes as a function of cellular context (depicted in UMAP embedding, left). (B) SURGE is a novel probabilistic model that uses matrix factorization to jointly learn a continuous representation of the cellular contexts defining each measurement (U), and the corresponding eQTL effect sizes specific to each learned context (V) based on observed expression (Y) and genotype (G) data. SURGE additional accounts for the effects of known covariates and sample repeat structure on gene expression (not shown in figure; see Methods). Assume there are N samples, T genome-wide independent variant-gene pairs, and K latent contexts. (C) Based on simulated data, we evaluated SURGE’s ability to reconstruct simulated latent contexts as measured by the average variance explained of the simulated latent contexts by the learned latent contexts (y-axis). We simulate 5 latent contexts and vary the sample size (x-axis) and the strength (variance; see Methods) of the interaction terms (colors). We fix the fraction of tests that are context-specific eQTLs for each context to .3 (see Methods). For each parameter setting, we run 10 independent simulations. Each dot is an independent simulation. (D). Based on simulated data, we evaluate SURGE’s ability to identify the number of simulated latent contexts across 10 independent simulations. The sample size was fixed to 250, the strength (variance) of the simulated interaction terms was fixed to .25, and the fraction of tests that are context-specific eQTLs for a particular context (see Methods) was fixed to .3. For each parameter setting, we run 10 independent simulations. Each dot is an independent simulation.

Indeed, eQTLs from adult bulk tissue samples fail to explain the majority of known disease loci [11,15–17]. It is therefore critical to identify eQTLs from diverse contexts in order to more fully characterize the molecular mechanisms underlying disease associated loci. Recent work has shown single-cell RNA-sequencing (scRNA-seq) provides unique data to uncover cell-type- and context-specific eQTLs; such higher-resolution data will naturally better reflect diverse cell types and cellular states, many of which would not be detectable from bulk RNA-seq [9,10,12–14,18,19].

However, the relevant contexts, such as cell type or state, that actually modulate genetic effects may not be known a priori. For example, genetic regulatory effects that are only present in a rare cell type, during intermediate stages of cellular differentiation [8,9] or in response to an environmental stimuli [7] that may not already be known to be disease relevant. Furthermore, an individual cell may be defined by multiple, overlapping contexts, such as both cell type and a perturbation response affecting partially overlapping sets of cells [9,20,21]. Contexts, such as differentiation progress or time, may manifest as continuous effects rather than discrete clusters. We developed SURGE (Single-cell Unsupervised Regulation of Gene Expression), a novel probabilistic model that uses matrix factorization to learn a continuous representation of the cellular contexts that modulate genetic effects. This includes the extent of relevance of each context to each cell or sample, and the corresponding eQTL effect sizes specific to each learned context, allowing for discovery of context-specific eQTLs without pre-specifying subsets of cells or samples.

First, we evaluate the statistical power of SURGE to identify latent contexts that modulate genetic effects on gene expression using simulated data. Next in a proof of concept experiment we apply SURGE to bulk gene expression measurements from ten GTEx version 8 tissues [5] to uncover the relevant contexts underlying eQTL regulatory patterns in bulk RNA-seq data. We then use SURGE to identify context-specific eQTLs in a single-cell data set consisting of 1.2 million peripheral blood mononuclear cells (PBMC) spanning 224 genotyped individuals [18]. Finally, we demonstrate the disease-relevance of SURGE context-specific eQTLs using colocalization analysis and stratified LD-score regression (S-LDSC) [22,23].

## Results

A standard approach to identify context-specific eQTLs is to quantify the effect of the interaction between genotype and a pre-specified cellular context on gene expression levels using a linear model (interaction-eQTLs) [8,13]. However, this approach requires pre-specifying which contexts, such as known cell types, to test for interaction, therefore inhibiting eQTL discovery in previously unstudied cellular contexts or uncharacterized cell types.

To address this issue, we developed SURGE, which uses a matrix factorization approach to uncover context-specific eQTLs without requiring pre-specification of the contexts of interest. SURGE achieves this goal by leveraging information across genome-wide variant-gene pairs to jointly learn both a continuous representation of the latent cellular contexts defining each measurement (henceforth referred to as SURGE latent contexts) and the interaction eQTL effect sizes corresponding to each SURGE latent context (Fig. 1B; see Methods). Importantly, SURGE allows for any individual measurement (such as a single cell) to be defined by multiple, overlapping contexts. From an alternative but equivalent lens, SURGE discovers the latent contexts whose linear interaction with genotype explains the most variation in gene expression levels. From this perspective, SURGE enables unsupervised discovery of the principal axes of genetic regulation of gene expression defining an eQTL data set. To increase power in detecting context-specific eQTLs, SURGE controls for the effects of known covariates as well as sample repeat structure induced by assaying multiple measurements (such as many cells) from the same individual on gene expression when identifying context-specific eQTLs (see Methods). Finally, SURGE automatically selects the number of relevant latent contexts by placing Automatic Relevance Determination priors distributions [24] on the inferred latent contexts (see Methods). The user simply would initialize the number of latent contexts to be large and greater than the likely number of underlying latent contexts present in the eQTL data set, and SURGE will prune unnecessary contexts during optimization (see Methods).

Recently, there have been two methods proposed to identify contexts related to genetic regulation of gene expression from eQTL data sets [25–27]. SURGE is unique from these methods in that it identifies contexts whose linear interaction with genotype explain the most variation in gene expression levels. Vochteloo et al. [25] identifies contexts such that the joint effect of the context and the interaction between the context and genotype maximize variation in residual gene expression. As such, this approach could identify contexts that have a linear effect on gene expression that is less related to genetic effects, which could happen in the common case where main effects of environment or cell type are larger than genetic interaction effects. Furthermore, this approach was not developed for application on single-cell eQTL data. Gewirtz et al. [26,27] propose a method to identify shared latent topics present in both expression and genotype data. Topics identified by this approach will not directly correspond to contexts whose interaction with genotype maximally explains variation in gene expression. Therefore, the goals of each method are distinct, and SURGE uniquely identifies contexts where interaction between genotype and context drive variation in gene expression.

We utilize a simulation framework to statistically quantify SURGE’s ability to accurately infer the latent contexts that alter genetic regulation of gene expression (see Methods). As expected, reconstruction of the simulated contexts depends on the sample size of the eQTL data set as well as the true effect size and number of context-specific eQTLs present in the simulated eQTL data set (Fig. 1C, Fig. S1). However, given a realistic eQTL data set containing 100 modest effect context-specific eQTLs (simulated realistic interaction variance 0.25 [8]; see methods) and sample size (n=250), SURGE accurately infers the simulated latent contexts (Fig. 1C, Fig. S1) as well as the number of simulated latent contexts (Fig. 1D, Fig. S2).

As a proof of concept in real sequencing data, we apply SURGE to model RNA-sequencing samples from 10 GTEx version 8 tissues (4169 individual-tissue pairs; Adrenal Gland, Colon-Sigmoid, Esophagus Mucosa, Muscle-Skeletal, Pituitary, Skin [not sun exposed suprapubic], Skin [sun exposed lower leg], Small Intestine terminal ileum, Stomach, and Thyroid), selected to be largely diverse with a small number of related tissues. SURGE identifies 8 latent contexts, all of which result in hundreds of genes with at least one SURGE interaction eQTL, or variant whose effect on expression changed with the SURGE latent context (eFDR <=.05, see Methods) (Fig. S3, Fig. S4, Table. S1). In this dataset, each RNA sample was extracted from a specific tissue, and while tissue identity information is not provided to SURGE, 6 of the 8 SURGE latent contexts capture differences in tissue type between the samples (Fig. 2A). SURGE latent context 1 (latent contexts ordered by PVE, see Methods), for example, isolates RNA samples from Muscle-Skeletal tissue; RNA samples derived from Muscle-Skeletal tissue have an average latent context 1 value of -1.82 (sdev 0.342), while RNA samples from other tissues have an average latent context 1 value of -0.011 (sdev 0.456). Furthermore, we discover SURGE latent context 4 and 7 cluster samples according to their known ancestry; samples from African Ancestry donors are strongly loaded on both latent context 4 and 7 (Fig. 2B, Fig. S5).

**Figure 2:**
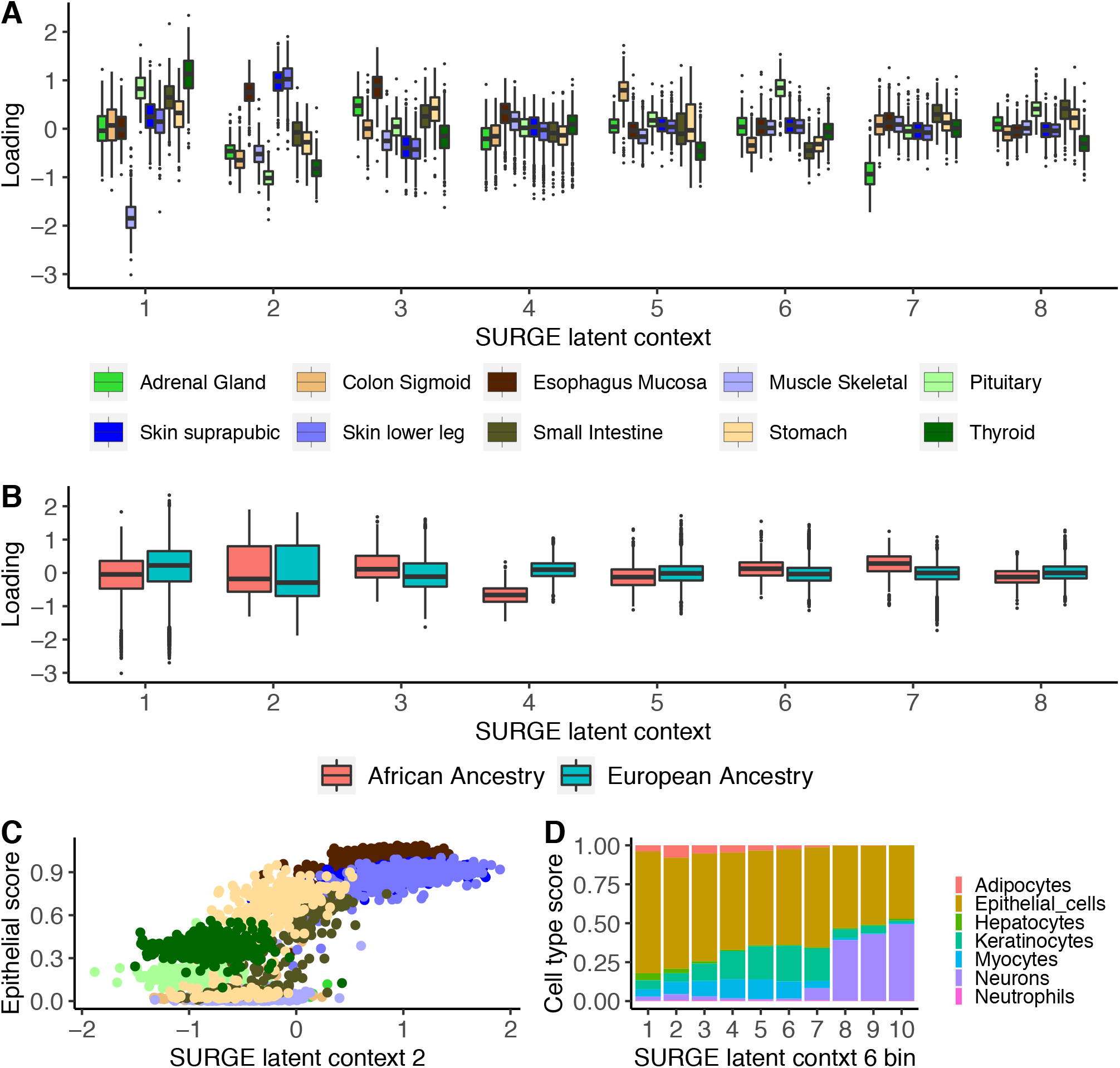
SURGE applied to GTEx v8 bulk RNA-seq samples. (A,B) SURGE latent context loadings of GTEx v8 RNA-seq samples (y-axis) stratified by (A) known tissue identity and (B) known ancestry for each of the 8 inferred SURGE latent contexts. (C) Scatter plot of SURGE latent context 2 loadings (x-axis) and xCell Epithelial cell type enrichment score (y-axis) for GTEx v8 RNA-seq samples colored by known tissue identity (same color palette as panel A). (D) GTEx v8 RNA-seq samples are separated into 10 quantile bins according to their value on SURGE latent context 6. The stacked bar plot depicts the average xCell cell-type enrichment scores across all samples normalized to sum to 1 (y-axis) in each of the 10 bins (x-axis).

Next, we intersect the learned SURGE latent contexts with previously computed computational estimates of each RNA sample’s cell type composition according to xCell (xCell infers cell type enrichment scores that reflect cell type composition based on external cell-type specific gene expression data) [28,29]. We find that the SURGE latent contexts are not simply identifying differences in tissue identity between the samples, but learning differences in cell type composition of samples both across tissues and within a single tissue (Fig. 2C, Fig. 2D, Fig. S6-S8). SURGE latent context 2, for example, is highly correlated with epithelial cell enrichment score across samples from all ten tissues (Fig. 2C). Moreover, many of the SURGE latent contexts capture complex multi-cell type composition continuums, not simply the change in proportions of a single cell type (Fig. 2D, Fig. S6). SURGE identifies latent contexts underlying cell type composition continuums even when applied to RNA samples from only a single tissue (see Methods, Fig. S9), demonstrating the importance of cell type composition differences across samples extracted from the same tissue. Importantly, we observe greater power to detect context-specific eQTLs with SURGE latent contexts than with the previously studied approach [28] of testing genetic interactions with cell type enrichment score estimates from xCell (see Methods, Fig. S10). In summary, SURGE identifies tissue-type, cell-type, and ancestry as the primary axes of genetic regulation of gene expression within GTEx eQTL data.

Next, we apply SURGE to a recently generated single cell eQTL data set consisting of 1.2 million PBMCs from 224 genotyped individuals [18]. 141 of these individuals have systemic lulus erythematosus (SLE) while the remainder are healthy. To mitigate the sparsity characteristic of 10X sequencing data, we aggregate cell level expression data across highly correlated cells to generate 22774 pseudocells (see Methods, Fig. S11) [21,30], aggregating on average 22 cells per pseudocell. Here, SURGE identifies 5 latent contexts, all of which resulted in hundreds of genes with at least one SURGE interaction eQTL (eFDR < .1, see Methods) (Fig. S12, Fig. S13, Table. S2).

The first 3 SURGE latent contexts capture continuous representations of distinct blood cell types while integrating biologically related cell types along a gradient within a single latent context (Fig. 3A, Fig. S14-S16). SURGE latent context 2, for example, is strongly loaded on Natural Killer (NK) cells, while still identifying fine-resolution differences distinguishing bright NK cells from dim NK cells (Fig. S14). Additionally, SURGE latent context 1 identifies subtle differences isolating monocytes derived from healthy individuals from monocytes derived from SLE individuals (Fig. S17; p < 2e-20, Wilcoxon rank sum test). Interestingly, SURGE latent context 4 and 5 do not capture broad expression trends related to cell types defining the top gene expression principal components (Fig. S18), and instead show strong correlation with expression of genes involved in specific biological processes (See Methods, Table S3, Table S4). For example, SURGE latent context 4 is correlated with genes that are extremely enriched in the Hallmark interferon-gamma response (odds ratio: 28.52, p < 4.2e-10) [31]. The interferon gamma response is a well-studied immune-related pathway shown to be involved in regulating SLE [18,32,33].

**Figure 3:**
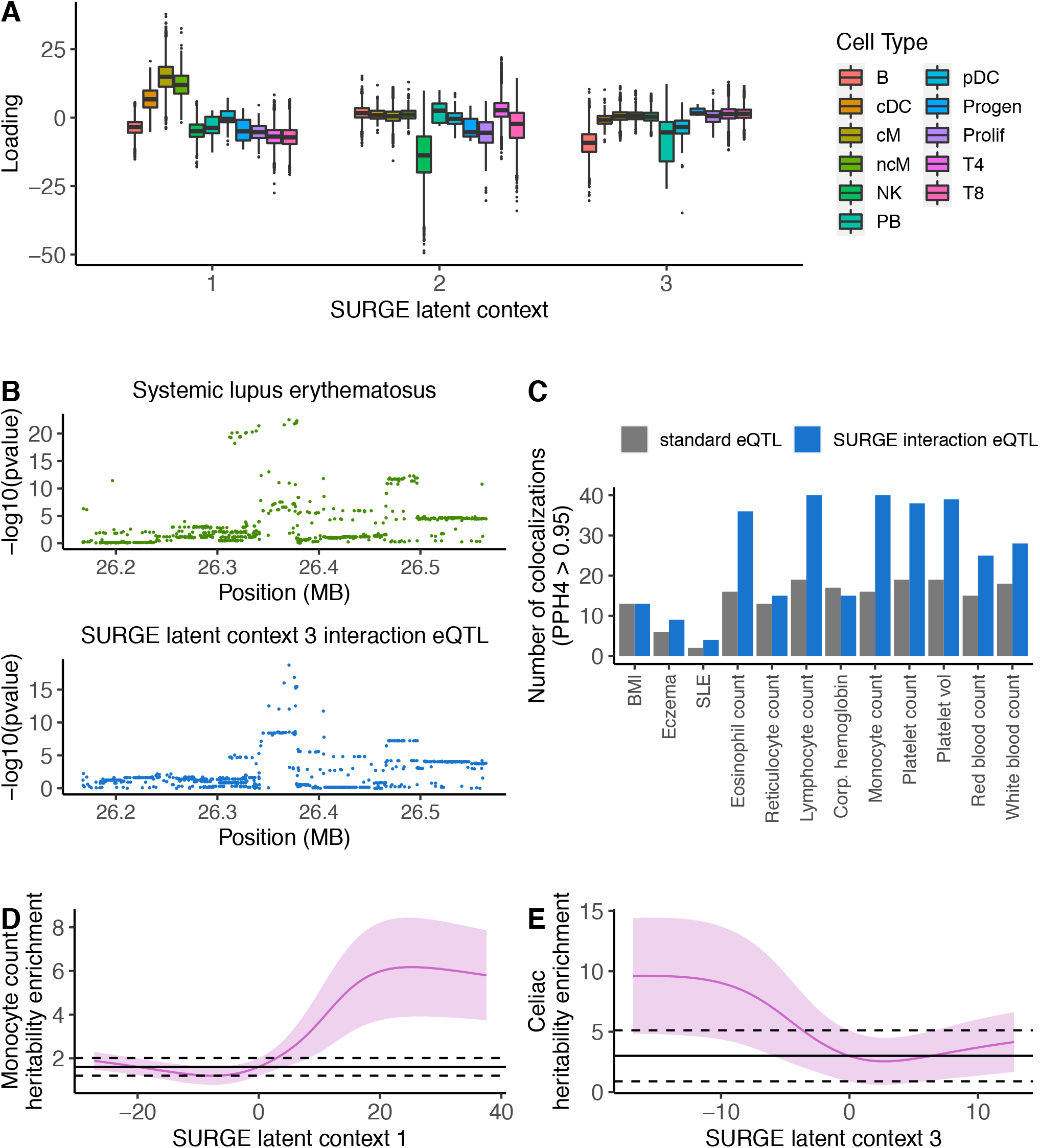
SURGE applied to PBMC single-cell eQTL data. (A) SURGE latent context loadings of pseudocells (y-axis) stratified by cell type (color) according to marker gene expression profiles for each of the top 3 SURGE latent contexts (x-axis). (B) Colocalization between SURGE latent context 3 interaction eQTL variant chr6:26370572:C:T for BTN3A2 and GWAS signal for SLE. (C) Number of colocalizations identified (PPH4 > .95; y-axis) between various GWAS studies (x-axis) and eQTLs identified from pseudocells. The number of colocalizations using standard eQTLs shown in grey and the number of unique colocalizations using SURGE interaction eQTLs, aggregated across contexts, shown in blue. (D,E) S-LDSC enrichment (y-axis) of squared standard eQTL effect sizes (black line) and SURGE predicted squared eQTL effect size at a specific SURGE latent context value (pink line at a specific x-axis position) within (D) monocyte count and (E) Celiac disease heritability. SURGE predicted eQTL effect sizes at a particular SURGE latent context value was calculated at 200 equally spaced positions along the range of SURGE latent context values. Black dashed line represents 95% confidence on the standard eQTL S-LDSC enrichment. Light pink region depicts 95% confidence on the SURGE predicted eQTL S-LDSC enrichment.

Finally, we evaluate the relationship between SURGE interaction eQTLs and disease-associated loci across diverse traits with genome-wide association studies (GWAS) available. Using coloc [22], we identify hundreds of colocalizations between SURGE interaction eQTLs and GWAS loci (Fig. 3B, Fig. 3C, Fig. S19). For example, a SURGE context 3 interaction eQTL for BTN3A2 colocalized with a GWAS signal for SLE (Fig. 3B). Furthermore, we identify significantly more trait colocalizations with SURGE interaction eQTLs relative to using standard eQTLs (Fig. 3C).

Next, we assess how eQTL enrichment in complex trait and disease heritability varied along the SURGE latent contexts using S-LDSC [23,34]. Briefly, we used SURGE to estimate eQTL effect sizes at multiple positions along each SURGE latent context, and then use S-LDSC to quantify the heritability enrichment of eQTLs identified at each position (see Methods). We note that this approach is not limited to SURGE interaction eQTLs, and could be applied to any analysis that infers interaction eQTLs. We observed that eQTL enrichment in complex trait and disease heritability significantly varies along the SURGE latent contexts for many diseases and complex traits (Fig. 3D, Fig. 3E, Fig. S20). For example, predicted eQTL effects in cells negatively loaded on SURGE latent context 3 (corresponding to a B-cell continuum) are approximately four times more enriched in Celiac disease heritability than static eQTLs (Fig. 3E). This result coincides with the previously-reported role of B-cells in Celiac disease [35,36]. Ultimately, this analysis highlights the importance of assessing eQTLs in disease-relevant contexts, as well as SURGE’s capacity for identifying disease-relevant contexts.

## Discussion

Here, we presented SURGE, a novel probabilistic model that identifies context-specific eQTLs from single-cell data without pre-specifying context, such as cell types or subsets of samples. SURGE leverages information from variant-gene pairs across the entire genome to learn a continuous representation of the cellular contexts defining each measurement, and the corresponding eQTL effect sizes specific to each learned context. Importantly, SURGE allows for unsupervised discovery of the principal axes of genetic regulation of gene expression within an eQTL data set, identifying cell-type, tissue-type, and ancestry when applied to GTEx tissue samples and highly resolved blood cell-types and gene programs when applied to blood-derived single cells. We demonstrated that eQTL enrichment in complex trait and disease heritability significantly varied along the SURGE latent contexts and ultimately, SURGE identified many trait-relevant loci that could not be detected through traditional eQTL approaches. Furthermore, large single-cell eQTL data sets are being rapidly generated containing cells spanning increasingly diverse cellular contexts. SURGE provides a statistically principled approach to uncover the dominant axes of genetic regulation of gene expression in such data.

## Methods

### SURGE model overview

The SURGE model is defined according to the following probability distributions:

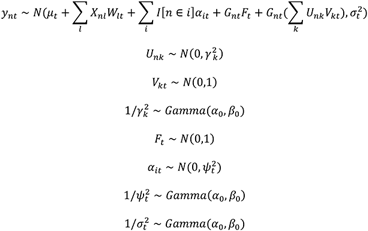

Here, *n* indexes RNA samples, *t* indexes independent variant-gene pairs being tested for eQTL analysis, and *i* indexes individuals. We use the notation *n* ∈ *i* to represent the instance where RNA sample *n* is drawn from the individual *i. y*_*nt*_ is the observed standardized gene expression (mean 0 and variance 1 for each test *t*) level of the gene corresponding to test *t* in sample *n*. We assume the gene expression data has been properly normalized prior to standardization. *G*_*nt*_ is the observed, standardized (described in more detail below) genotype of the variant corresponding to test *t* in sample *n. X*_*nl*_ is the observed value of covariate *l* for sample *n* . SURGE infers the values of:

- *F*_*t*_: the eQTL effect size of test *t* that is shared across samples
- *V*_*kt*_: the eQTL effect size of test *t* for latent context *k*
- *U*_*nk*_: the latent context value of sample *n* on factor *k*
- *μ*_*t*_: the intercept of each test
- *W*_*lt*_: The effect size of covariate *l* on the gene corresponding to test *t*
- *α*_*it*_: the random effect intercept for each individual for each test
- 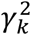: The variance of the values in latent context *k*
- 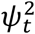: The variance of intercept corresponding to each individual in test *t*
- 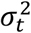: The residual variance in gene expression levels in test *t*

*a*_0_, and *β*_0_ are model hyper-parameters set to provide non-informative priors while stabilizing optimization. In practice we set *α*_0_ to 1e^-3^ and *β*_0_ to 1e^-3^.

To standardize the genotype of the variant corresponding to test *t*, we center the genotype vector to have mean 0 across samples and then we scale the genotype vector for test *t* (*G*_\**t*_) by the standard deviation of *Y*_\**t*_*/G*_\**t*_. This scaling encourages the low-dimensional factorization (*UV*) to explain variance equally across tests instead of preferentially explaining variance in tests with small variance in *Y*_\**t*_*/G*_\**t*_.

It is worth highlighting that a mean-zero gaussian prior is placed on *U*_*nk*_ in order to produce interpretable assignments of samples to factors. The level of regularization of that prior is learned separately for each latent context 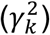, allowing SURGE to zero-out (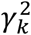 approaches 0) irrelevant contexts and automatically learn the effective number of latent contexts. This approach has been used by others for inference of the number of effective components in more traditional matrix factorizations [37,38] and is similar to Automatic Relevance Determination [24].

### SURGE optimization

We approximate the posterior distribution of all latent variables 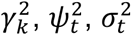 using mean-field variational inference [39]. The goal of variational inference is to minimize the KL-divergence between *q*(*Z*) and *p*(*Z*|*Y, G, X*), which can be written as *KL*(*q*(*Z*)||*p*(*Z*|*Y, G, X*). Here, *q*(*Z*) is a simple, tractable distribution that is used to approximate *p*(*Z*|*Y, G, X*). We used the “mean-field approximation” for *q*(*Z*) such that all latent variables are independent of one another. More specifically:

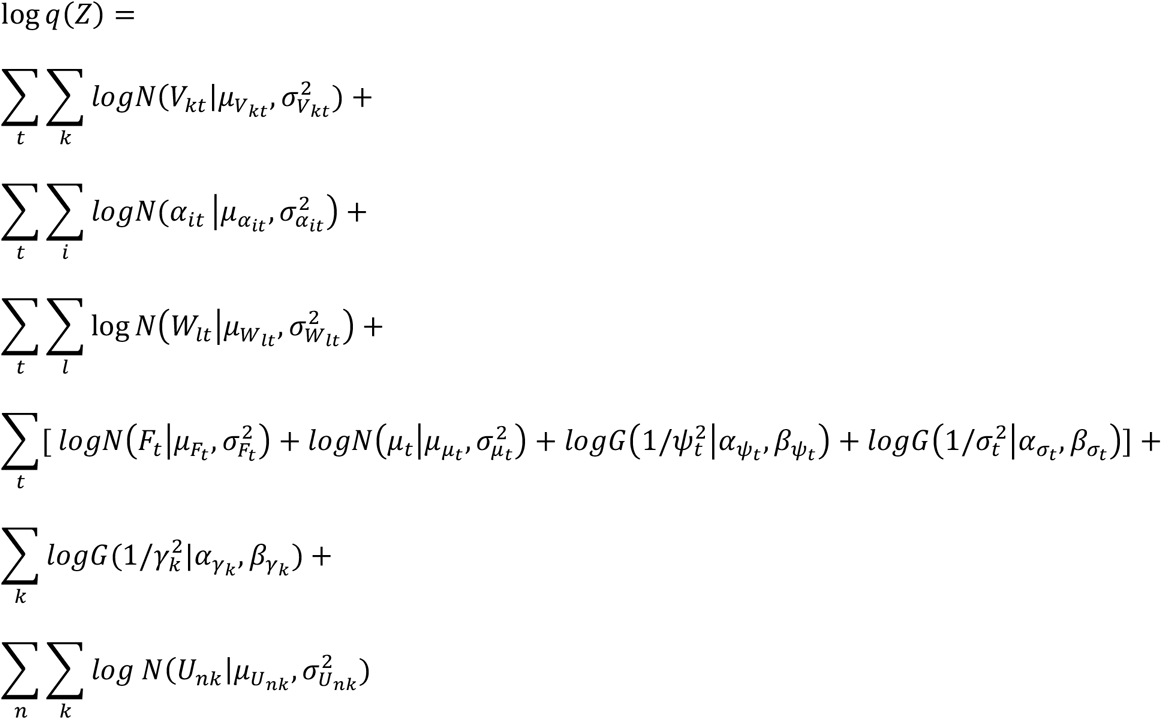

Where *N*(*x*|*μ, σ*^2^) is a univariate normal distribution parameterized by mean *μ* and variance *σ*^2^ and *G*(*X*|*α, β*) is a univariate gamma distribution parameterized by *α* and *β*. It can be shown that minimizing the KL-divergence *KL*(*q*(*Z*)||*p*(*Z*|*Y, G, X*) is equivalent to maximizing the evidence lower bound (ELBO):

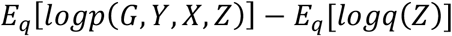

Therefore, we will frame SURGE optimization from the perspective of maximizing the ELBO with respect to the parameters defining *q*(*Z*), or the variational parameters. Noteworthy is *p*(*G, Y, X, Z*) is explicitly defined in the methods section “SURGE model overview” and can be easily computed. The approach we take to maximize the ELBO is through coordinate ascent [39], iteratively updating the variational distribution each latent variable, while holding the variational distributions of all other latent variables fixed. Accordingly, the ELBO is guaranteed to monotonically increase after each variational update. In the case of the SURGE model, each update is available in closed form (see Supplement).

Optimization of variational parameters is performed as follows: we randomly initialize all variational parameters (see below section entitled “Random initialization for SURGE optimization”) and then iteratively loop through all latent variables in *Z* and update the variational parameters corresponding to that latent variable until we reach convergence.

To assess convergence, we assess the change in ELBO from one iteration to the next. We consider the model converged when the change in ELBO is less than 1e-2.

### Random initialization for SURGE optimization

It is important to note that mean-field variational inference is not guaranteed to converge to the global optima of the ELBO. To mitigate the effects of local optima, we recommend optimizing multiple models with different random initializations and using the parameters learned from the model that achieves the largest ELBO.

### Percent variance explained of SURGE latent contexts

Following the approach taken by [40], we define the “Percentage Variance Explained” (PVE) of the *k*^*th*^ latent context as:

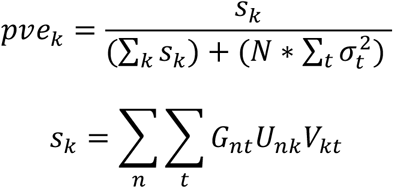

As stated in [40], this approach is a measure of the amount of signal in data set that is identified by the *k*^*th*^ latent context. However, the name “percentage variance explained” should be considered loosely as the factors are not orthogonal.

### Removing irrelevant latent contexts

Upon model convergence, we remove latent contexts with PVE ≤ 1*e*^−5^.

### Simulation experiments

To assess SURGE’s ability to accurately capture contexts underlying context-specific eQTLs we performed the following simulation experiment:

We randomly generated genotype and expression matrices across 1000 variant-gene pairs and *N* RNA samples. For each simulated variant-gene pair, we simulated the genotype vector (*G*) across the *N* samples according to the following probability distributions:

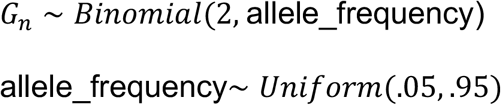

Then, we simulated the expression vector (*y*) across the *N* samples using that variant-gene pair’s simulated genotype vector according to the following probability distributions:

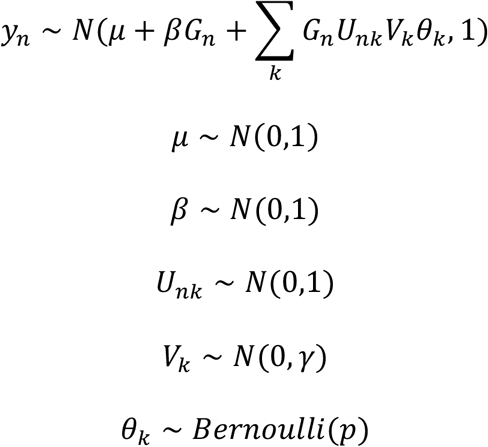

Using this simulation, we can evaluate SURGE’s ability to re-capture the simulated latent contexts (U) (Fig. 1C, Fig. S1) as a function of the simulation hyper-parameters:

- The number of latent contexts (K)
- The sample size (*N*)
- The strength of the interaction terms (*γ*)
- The fraction of tests that are context-specific eQTLs for a particular context (*p*)

We can also access SURGE’s ability to accurately estimates the number of relevant contexts (K) (Fig 1D, Fig. S2) by only retaining contexts with PVE > 1*e*^−5^.

### Selection of variant-gene pairs used for optimization

SURGE optimization (ie. learning the SURGE latent contexts) requires an input expression matrix and genotype matrix. As specified above, both matrices should be of dimension *N*X*T*, where *N* is the number of RNA samples and *T* is the number of genome-wide independent variant gene pairs. We desire each variant-gene pair to be independent of one another because we want the SURGE to infer eQTL patterns that are persistent across the genome, not specific to a single gene or variant.

To encourage the expression and genotype data to consist of independent variant-gene pairs we limit there to be a single variant-gene pair selected for each gene and limit there to be a single variant-gene pair selected for each variant.

Furthermore, it has been shown that context-specific eQTLs are more likely to be standard eQTLs than not. We therefor limit variant-gene pairs used for SURGE optimization to those that are standard eQTLs within the data set (more details presented below). For computational efficiency, we recommend using a maximum of 2000 genome-wide independent variant-gene pairs for SURGE optimization.

### SURGE interaction-eQTLs

SURGE optimization on a subset of genome-wide independent variant-gene pairs will result in approximations to the posterior distributions of the SURGE latent contexts (U) as well as eQTL effect sizes for each of the SURGE latent contexts for only the genome-wide independent variant gene pairs (V). It is of interest, however, to call interaction eQTLs with respect to each of the SURGE latent contexts for *all* variant gene-pairs, not just the subset of variant-gene pairs that are genome-wide independent and used for SURGE optimization.

Therefore, to identify SURGE interaction-eQTL for an arbitrary variant-gene pair we treat the expected value of the inferred posterior distribution on the SURGE latent contexts (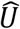: dim NXK) as observed and optimize the following linear mixed model for each variant-gene pair. The linear mixed model is as follows:

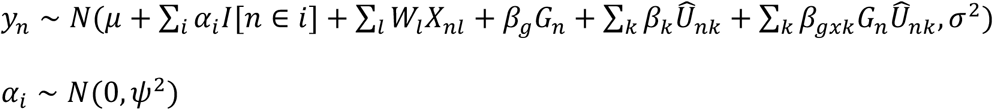

Here:

- *y*_*n*_ is the observed expression level of the gene corresponding to the variant-gene pair in sample *n*
- *g*_*n*_ is the observed genotype of the variant corresponding to the variant-gene pair in sample *n*
- *X*_*nl*_ is the observed value of covariate *l* in sample *n*
- *μ* is the intercept
- *α*_*i*_ is the random effect intercept for individual *i*. We use the notation *n* ∈ *i* to represent the case where sample *n* is drawn from individual *i*
- *W*_*l*_ is the fixed effect for covariate *l*
- *β*_*g*_ is the fixed effect for genotype
- *β*_*k*_ is the fixed effect of the *k*^*th*^ latent context
- *β*_*gxk*_ is the fixed effect of the interaction between the *k*^*th*^ latent context and genotype

We use the R package ‘lme4’ to quantify the significance of all K interaction terms: *β*_*gx*1_, …, *β*_*gxk*_, …, *β*_*gxk*_. Intuitively, if the *k*^*th*^ interaction term (*β*_*gxk*_) is significant, it implies that the eQTL effect size of this variant-pairs significantly changes along latent context *k*.

### Calibration of SURGE interaction eQTLs using permutation analysis

P-values resulting from SURGE interaction-eQTL analysis are potentially inflated due to SURGE interaction eQTLs being identified from the same data used to learn the SURGE latent contexts. This statistical phenomenon is known as “double-dipping” and there exist well-studied approaches to ensure statistical calibration in the presence of “double-dipping” [41–44]. We use a conservative, permutation analysis to generate an empirical null distribution of gene-level p-values that can be used to calibrate the observed gene-level p-values. The permutation analysis consisted of (1) permuting genotype of each individual, (2) re-optimizing SURGE latent contexts (U) using the permuted genotype data, and (3) calling SURGE interaction-eQTLs with the permuted genotype data and the SURGE latent contexts learned using the permuted genotype data. More specifically, we generated a single permutation of genotype that was used across all variants to ensure we did not break the correlation structure across variant-gene pairs. In addition, we only permuted genotype across individuals, not RNA samples, to ensure we preserved sample repeat structure expression effects. This means that multiple RNA samples from the same individual will always have the same genotype values in the permutation run. Lastly, similar to previous permutation experiments on linear-interaction effects [8,45], we only permuted the genotype variable in the interaction term while leaving the fixed effect of genotype un-permuted (for both SURGE optimization and SURGE interaction-eQTL calling).

Given both the observed and permuted SURGE interaction eQTL p-values, we generated gene-level p-values using Bonferonni correction for both observed and permuted interaction eQTLs. We then evaluated genome-wide significance of the observed gene-level p-values using empirical FDR (eFDR) [46] calibrated with the permuted gene-level p-values. This approach was performed independently for each SURGE latent context.

In the real (un-permuted) data, we only called SURGE interaction-eQTLs for SURGE latent contexts with PVE > 1e^-5^. Unsurprisingly, permuted SURGE latent contexts consistently explained less PVE than un-permuted SURGE latent contexts (Fig. S3, Fig. S12); there existed zero permuted SURGE latent contexts explaining PVE > 1e^-5^ across all experiments. Therefore, if Z SURGE latent contexts have PVE > 1e^-5^, we selected the top Z permuted SURGE latent contexts to be used in the permuted SURGE interaction eQTL analysis.

### Application of SURGE to GTEx samples from 10 tissues: expression quantification

To normalize expression from samples from 10 GTEx tissues (Adrenal gland, Colon-sigmoid, Esophagus-Mucosa, Muscle-Skeletal, Pituitary, Skin-not-sun-exposed, Skin-sun-exposed, small-intestine-terminal-ileum, Stomach, Thyroid), we concatenated log-TPM expression measurements across all samples used in the GTEx v8 eQTL analysis for one of those tissues [5]. We also limited to genes that were tested for eQTLs in the GTEx v8 analysis [5] in all 10 tissues. Next, we quantile normalized this matrix to ensure each sample had an equivalent distribution across genes and then standardized each gene (mean 0 and standard deviation 1). We excluded RNA samples that were outliers (Z-score >= 4) according to Mahalanobis distance computed on 80 expression PCs.

### Application of SURGE to GTEx samples from 10 tissues: standard eQTL calling

We first tested for standard eQTLs, or association between genotype and the concatenated (across tissues) expression vector described above in “Application of SURGE to GTEx samples from 10 tissues: expression quantification”. For this analysis, we limited to genes that passed filters described in “Application of SURGE to GTEx samples from 10 tissues: expression quantification”. We then limited to variants with MAF >= .05 that were less than 50KB from the transcription start site of a gene. We controlled for the effects of 80 expression PCs and 4 genotype PCs (as recommended by [5] given the sample size). We assessed genome-wide significance according to a gene-level Bonferonni correction, followed by a genome-wide Benjamini-Hochberg correction.

### Application of SURGE to GTEx samples from 10 tissues: SURGE optimization

To select a subset of variant-gene pairs to be used for SURGE model optimization, we first limited to variant-gene pairs that were standard eQTLs (FDR <= .05; see “Application of SURGE to GTEx samples from 10 tissues: standard eQTL calling”). This was done to ensure a higher fraction of the variant-gene pairs used for SURGE optimization were context-specific eQTLs as it is known standard eQTLs are more likely to be context-specific eQTLs than variant-gene pairs that are not standard eQTLs. Furthermore, we limited to the most significant variant per gene amongst the 2000 most significant genes and removed a variant-gene pair if the variant was already in the training set for its association with a more significant gene. This yielded 1,996 genome-wide independent variant-gene pairs used for SURGE optimization. We then ran SURGE under default parameter settings over these genome-wide independent variant-gene pairs. We included 80 expression PCs and 4 genotype PCs as covariates in SURGE optimization. The converged SURGE model resulted in 8 latent contexts with PVE > 1*e*^−5^ and hundreds of genome-wide significant SURGE interaction eQTLs (eFDR <= .05) (Fig. S4, Table S1).

### Application of SURGE to GTEx samples from a single tissue

To run SURGE on GTEx samples from a single GTEx tissue, we took a very similar approach to that described in “Application of SURGE to GTEx samples from 10 tissues: expression quantification”, “Application of SURGE to GTEx samples from 10 tissues: standard eQTL calling”, and “Application of SURGE to GTEx samples from 10 tissues: SURGE optimization”. The only difference is that we now limit to samples from the tissue of interest. Furthermore, we now only control for 60 expression PCs and 2 genotype PCs during standard eQTL calling and SURGE optimization. The converged model resulted in 1 latent contexts with PVE > 1*e*^−5^ and 1287 genome-wide significant SURGE interaction eQTLs (eFDR <= .05).

### Application of SURGE to PBMC single cell eQTL data: pseudocell expression quantification

We imported raw, un-normalized UMI counts from [18]. We used SCRAN [47] to generate log-normalized counts for each cell. We removed genes that were expressed in fewer than .5% of cells. We then limited to the top 6000 highly variable genes via the Scanpy function “highly_variable_genes” [48]. We then removed the effects of sequencing batch using Combat [49] as implemented in Scanpy. We then scaled each gene to have mean 0 and variance 1, with a maximum absolute value of 10 to mitigate outlier effects as implemented by “scanpy.pp.scale”.

Next, we sought to generate pseudocells that represented groupings of highly correlated cells within an individual. We first removed individuals from this analysis with fewer than 2500 cells. Next we performed Leiden clustering as implemented by Scanpy [50] independently in each individual using all default parameters, except we used a fine-grained cluster resolution of 10. Here, each leiden cluster corresponds to a pseudocell. We took the average expression across all cells assigned to the pseudocell to estimate the expression profile of the pseudocell. Finally, we standardized each gene (across pseudocells) to have mean 0 and standard deviation 1, again capping the absolute value of standardized scores to be 10 to mitigate outlier effects. We excluded RNA pseudocells that were outliers (Z-score >= 4) according to Mahalanobis distance computed on 30 expression PCs.

### Application of SURGE to PBMC single cell eQTL data: standard eQTL calling

We first tested for standard eQTLs, or association between genotype and the expression vector across pseudocells described above in “Application of SURGE to Ye-lab generated single cell eQTL data: pseudocell expression quantification”. For this analysis, we limited to genes that passed filters described in “Application of SURGE to Ye-lab generated single cell eQTL data: pseudocell expression quantification”. We then limited to variants with MAF >= .05 that were less than 200KB from the transcription start site of a gene. We controlled for the effects of 30 expression PCs and 2 genotype PCs. We controlled for sample-repeat structure stemming from multiple pseudocells originating from the same individual using a random effects intercept for each individual. We assessed genome-wide significance according to a gene-level Bonferonni correction, followed by a genome-wide Benjamini-Hochberg correction.

### Application of SURGE to PBMC single cell eQTL data: SURGE optimization

To select a subset of variant-gene pairs to be used for SURGE model optimization, we first limited to variant-gene pairs that were standard eQTLs (FDR <= .05; see “Application of SURGE to Ye-lab generated single cell eQTL data: standard eQTL calling”). This was done to ensure a higher fraction of the variant-gene pairs used for SURGE optimization were context-specific eQTLs as it is known standard eQTLs are more likely to be context-specific eQTLs than variant-gene pairs that are not standard eQTLs. Furthermore, we limited to the most significant variant per gene amongst the 2000 most significant genes and removed a variant-gene pair if the variant was already in the training set for its association with a more significant gene. We than ran SURGE under default parameter settings over these genome-wide independent variant-gene pairs. We included 30 expression PCs and 2 genotype PCs as covariates in SURGE as well as a random effect intercept term for each individual. The converged SURGE model resulted in 5 latent contexts with PVE > 1*e*^−5^ and hundreds of genome-wide significant SURGE interaction eQTLs (eFDR <= .1) (Fig. S13, Table S2).

### Gene set enrichment analysis

We tested enrichment of genes whose expression levels was highly correlated with SURGE latent contexts (identified when SURGE was applied to single-cell PBMC data) within known gene sets. Specifically for each SURGE latent context, we identified the 50 genes whose expression levels across pseudocells were most strongly correlated (absolute value of correlation coefficient) with the SURGE latent context. We then tested gene set enrichment of these 50 genes relative to all genes that passed filters described in Methods section “Application of SURGE to PBMC single cell eQTL data: pseudocell expression quantification”. We tested enrichment of these strongly correlated genes in both the Hallmark gene set and the MSigDB Biological Process gene set [31] (Table S3, Table S4).

### Application of Stratified LD Score Regression (S-LDSC)

Recall, SURGE interaction eQTLs for a specific variant-gene pair can be identified by evaluating the following likelihood (see Methods section “Surge interaction eQTLs” for more details):

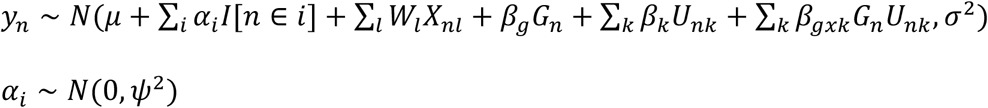

Upon maximizing this likelihood (assume 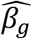 and 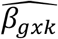 are the estimated values of *β*_*g*_ and *β*_*gxk*_ that maximize the likelihood), we can estimate the expected eQTL effect size for the variant-gene pair for a particular value of a latent context of U using the following function:

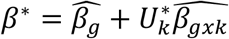

Here, *β*^*^ is the expected eQTL effect size for the particular variant-gene pair when the *k*^*th*^ latent context value of *U* is equal to 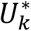. Ultimately, this enables us to compute the expected eQTL effect size for all variant-gene pairs when the *k*^*th*^ latent context value of U is equal to 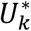.

We use the above expectation to assess how eQTL enrichment in complex trait and disease heritability varied along the SURGE latent contexts. Specifically, for each SURGE latent context we generated 200 equally-spaced positions along the range of SURGE latent context values. For each of those 200 positions, we computed the expected eQTL effect sizes (using the above expectation of *β*^*^) for all variant-gene pairs. We then used the squared expected eQTL effect size across variant-gene pairs as annotation in S-LDSC [23,34] along with all BaselineLD v2.2 annotations excluding four QTL related annotations (“GTEx_eQTL_MaxCPP”, “BLUEPRINT_H3K27acQTL_MaxCPP”, “BLUEPRINT_H3K4me1QTL_MaxCPP”, “BLUEPRINT_DNA_methylation_MaxCPP”). If a given variant mapped to multiple genes, we used the sum of squared expected eQTL effect sizes across genes as the annotation similar to [16]. This analysis was done for each of the 200 equally spaced positions for each of the 5 SURGE latent contexts identified when SURGE was run on the single cell PBMC eQTL data.

## Declarations

### Ethics approval and consent to participate

Not applicable.

### Consent for publication

Not applicable.

### Availability of data and materials

SURGE software is available on github at https://github.com/BennyStrobes/surge. The GTEx v8 data [5] can be downloaded from the dbGaP website under phs000424.v8.p2 and on the GTEx portal (http://gtexportal.org/). PBMC single-cell eQTL [18] expression data are available in the Human Cell Atlas Data Coordination Platform and at GEO accession number GSE174188. PBMC single-cell eQTL [18] genotype data are available at dbGap accession number phs002812.v1.p1

### Authors’ contributions

AB and BJS proposed the idea for the project. BJS developed the SURGE model and performed the analysis. BJS and AB wrote the manuscript. KT aided in deriving the variational updates for SURGE. JP provided review of SURGE code. JP, GQ, and AB suggested relevant downstream analysis to run based on SURGE output. MGG, RP, and CJY provided guidance on analysis of single-cell eQTL data.

## Acknowledgements

We thank Alkes L. Price for insights into the S-LDSC analysis and for providing comments on the manuscript.

