## Supplementary Note for "Uncovering context-specific genetic-regulation of gene expression from single-cell RNA-sequencing using latent-factor models"

**SURGE model overview**

The SURGE model is defined according to the following probability distributions:

$$y_{nt} \sim N(\mu_t + \sum_l X_{nl}W_{lt} + \sum_i I[n \in i]\alpha_{it} + G_{nt}F_t + G_{nt}(\sum_k U_{nk}V_{kt}), \sigma_t^2)$$

$$U_{nk} \sim N(0, \gamma_k^2)$$

$$V_{kt} \sim N(0,1)$$

$$1/\gamma_k^2 \sim Gamma(\alpha_0, \beta_0)$$

$$F_t \sim N(0,1)$$

$$\alpha_{it} \sim N(0, \psi_t^2)$$

$$1/\psi_t^2 \sim Gamma(\alpha_0, \beta_0)$$

$$1/\sigma_t^2 \sim Gamma(\alpha_0, \beta_0)$$

Here, $n$ indexes RNA samples, $t$ indexes independent variant-gene pairs being tested for eQTL analysis, and $i$ indexes individuals. We use the notation $n \in i$ to represent the instance where RNA sample $n$ is drawn from the individual $i$. $y_{nt}$ is the observed normalized gene expression (mean 0 and variance 1 for each test $t$) level of the gene corresponding to test $t$ in sample $n$. $G_{nt}$ is the observed, standardized genotype of the variant corresponding to test $t$ in sample $n$. $X_{nl}$ is the observed value of covariate $l$ for sample $n$ .

To standardize the genotype of the variant corresponding to test $t$, we center the genotype vector to have mean 0 across samples and then we scale the genotype vector for test $t$ ($G_{*t}$) by the standard deviation of $Y_{*t}/G_{*t}$. This scaling encourages the lowdimensional factorization ($UV$) to explain variance equally across tests instead of preferentially explaining variance in tests with small variance in $Y_{*t}/G_{*t}$.

SURGE infers the values of:

- $F_t$: the eQTL effect size of test $t$ that is shared across samples
- $V_{kt}$: the eQTL effect size of test $t$ for latent context $k$
- $U_{nk}$: the latent context value of sample $n$ on factor $k$
- $\mu_t$: the intercept of each test
- $W_{lt}$: The effect size of covariate $l$ on the gene corresponding to test $t$
- $\alpha_{it}$: the random effect intercept for each individual for each test
- $\gamma_k^2$: The variance of the values in latent context $k$
- $\psi_t^2$: The variance of intercept corresponding to each individual in test $t$
- $\sigma_t^2$: The residual variance in gene expression levels in test $t$

$a_0$, and $\beta_0$ are model hyper-parameters set to provide non-informative priors while stabilizing optimization. In practice we set $\alpha_0$ to 1e$^{-3}$ and $\beta_0$ to 1e$^{-3}$.

**SURGE inference overview**

We approximate the posterior distribution of all latent variables [$Z$ =($F_t$, $V_{kt}$, $U_{nk}$, $\mu_t$, $W_{lt}$, $\alpha_{it}$, $\gamma_k^2$, $\psi_t^2$, $\sigma_t^2$)] using mean-field variational inference. Variational inference seeks to minimize the KL-divergence from an approximating distribution $q(Z)$ to the exact posterior $p(Z|Y, G, X)$. We used the "mean-field approximation" for $q(Z)$ such that all latent variables are independent of one another. More specifically:

$\log q(Z) =$

$$\sum_t \sum_k logN(V_{kt}|\mu_{V_{kt}}, \sigma^2_{V_{kt}}) +$$

$$\sum_t \sum_i logN(\alpha_{it}\,|\mu_{\alpha_{it}}, \sigma^2_{\alpha_{it}}) +$$

$$\sum_t \sum_l \log N\left(W_{lt}\middle|\mu_{W_{lt}}, \sigma^2_{W_{lt}}\right) +$$

$$\sum_t [\, logN\left(F_t\middle|\mu_{F_t}, \sigma^2_{F_t}\right) + logN\left(\mu_t\middle|\mu_{\mu_t}, \sigma^2_{\mu_t}\right) + logG\left(1/\psi^2_t\middle|\alpha_{\psi_t}, \beta_{\psi_t}\right) + logG\left(1/\sigma^2_t\middle|\alpha_{\sigma_t}, \beta_{\sigma_t}\right)] +$$

$$\sum_k logG(1/\gamma^2_k|\alpha_{\gamma_k}, \beta_{\gamma_k}) +$$

$$\sum_n \sum_k log\ N(U_{nk}|\mu_{U_{nk}}, \sigma^2_{U_{nk}})$$

Where $N(x|\mu, \sigma^2)$ is a univariate normal distribution parameterized by mean $\mu$ and variance $\sigma^2$ and $G(X|\alpha, \beta)$ is a univariate gamma distribution parameterized by $\alpha$ and $\beta$.

It can be shown that minimizing the KL-divergence $KL(q(Z)||p(Z|Y, G, X)$ is equivalent to maximizing the evidence lower bound (ELBO):

$$E_q[logp(G, Y, X, Z)] - E_q[logq(Z)]$$

The approach we take to maximize the ELBO is through coordinate ascent, iteratively updating the variational distribution each latent variable, while holding the variational distributions of all other latent variables fixed. Accordingly, the ELBO is guaranteed to monotonically increase after each variational update. In the case of the SURGE model, each update is available in closed form (shown below).

**SURGE coordinate ascent variational inference update equations**

Below we give the closed form update for each latent variable in SURGE, which is applied at each iteration of the coordinate ascent variational inference algorithm. We use the notation $\langle z \rangle$ to represent the expected value of the random value z with respect to the variational distribution ($\langle z \rangle = E_q[z]$)

Latent contexts ($U_{nk}$)

For each sample $n$ and latent context $k$,

Prior distribution: $p(U_{nk}) \sim N(0, \gamma_k^2)$

Variational distribution: $q(U_{nk}) \sim N(\mu_{U_{nk}}, \sigma^2_{U_{nk}})$

where the updates are:

$$\sigma^2_{U_{nk}} = \left( \left( \sum_t \frac{1}{\langle \sigma_t^2 \rangle} G_{nt}^2 \langle V_{kt}^2 \rangle \right) + \frac{1.0}{\gamma_k^2} \right)^{-1}$$

$$\mu_{U_{nk}} = \sigma^2_{U_{nk}} \sum_t \frac{1.0}{\langle \sigma_t^2 \rangle} G_{nt} \langle V_{kt} \rangle \left( r_{nt}^{U_{nk}} \right)$$

$$r_{nt}^{U_{nk}} = Y_{nt} - \langle \mu_t \rangle - \sum_l X_{nl} \langle W_{lt} \rangle - \sum_i I[n \in i] \langle \alpha_{it} \rangle - G_{nt} \langle F_t \rangle - G\_nt \sum_{j!=k} \langle U_{nj} \rangle \langle V_{jt} \rangle$$

Latent contexts eQTL effect sizes ($V_{kt}$)

For each test $t$ and latent context $k$

Prior distribution: $p(V_{kt}) \sim N(0, 1)$

Variational distribution $q(V_{kt}) \sim N(\mu_{V_{kt}}, \sigma^2_{V_{kt}})$

where the updates are:

$$\sigma_{V\,kt}^2 = \left(\frac{1.0}{\langle\sigma_t^2\rangle}\sum_n G_{nt}^2\langle U_{nk}^2\rangle + 1\right)^{-1}$$

$$\mu_{V\,kt} = \sigma_{V\,kt}^2 \sum_n \frac{1.0}{\langle\sigma_t^2\rangle} G_{nt}\langle U_{nk}\rangle\left(r_{nt}^{V_{kt}}\right)$$

$$r_{nt}^{V_{kt}} = Y_{nt} - \langle\mu_t\rangle - \sum_l X_{nl}\langle W_{lt}\rangle - \sum_i I[n \in i]\langle\alpha_{it}\rangle - G_{nt}\langle F_t\rangle - \ G_{nt}\sum_{j!=k}\langle U_{nj}\rangle\langle V_{jt}\rangle$$

Shared eQTL effect sizes ($F_t$)

For each test *t*

Prior distribution: $p(F_t) \sim N(0,1)$

Variational distribution $q(F_t) \sim N(\mu_{F\,t}, \sigma_{F_t}^2)$

Where updates are:

$$\sigma_{F\,t}^2 \ = \left(\frac{1.0}{\langle\sigma_t^2\rangle}\sum_n G_{nt}^2 + 1\right)^{-1}$$

$$\mu_{F\,t} = \sigma_{F_t}^2 \sum_n \frac{1}{\langle\sigma_t^2\rangle} G_{nt}\left(r_{nt}^{F_t}\right)$$

$$r_{nt}^{F_t} \ = Y_{nt} - \langle\mu_t\rangle - \sum_l X_{nl}\langle W_{lt}\rangle - \sum_i I[n \in i]\langle\alpha_{it}\rangle - \ G_{nt}\sum_k\langle U_{nk}\rangle\langle V_{kt}\rangle$$

Effects of known covariates on gene expression ($W_{lt}$)

For each covariate *l* and each test *t*

Prior distribution: $p(W_{lt}) \sim N(0,M)$

Note M is picked to be infinitely large such that $\frac{1}{M} = 0$. This provides no regularization on the effects of covariates and allows as much expression variation as possible to be captured by covariates, leading to conservative estimates of genetic effects on gene expression.

Variational distribution $q(W_{lt}) \sim N(\mu_{W_{lt}}, \sigma^2_{W_{lt}})$

Where updates are:

$$\sigma^2_{W_{lt}} = \left(\frac{1.0}{\langle\sigma_t^2\rangle}\sum_n X_{nl}^2\right)^{-1}$$

$$\mu_{W_{lt}} = \sigma^2_{W_{lt}}\sum_n \frac{1}{\langle\sigma_t^2\rangle} X_{nl}(r_{nt}^{W_{lt}})$$

$$r_{nt}^{W_{lt}} = Y_{nt} - \langle\mu_t\rangle - \sum_{j!=l} X_{nj}\langle W_{jt}\rangle - \sum_i I[n \in i]\langle\alpha_{it}\rangle - G_{nt}\langle F_t\rangle - \ G_{nt}\sum_k \langle U_{nk}\rangle\langle V_{kt}\rangle$$

<u>Expression intercept ($\mu_t$)</u>

For each test *t*

Prior distribution: $p(\mu_t) \sim N(0, M)$

Note M is picked to be infinitely large such that $\frac{1}{M} = 0$. This provides no regularization on the intercept and allows as much expression variation as possible to be captured by the intercept, leading to conservative estimates of genetic effects on gene expression.

Variational distribution $q(\mu_t) \sim N(\mu_{\mu_t}, \sigma^2_{\mu_t})$

Where updates are:

$$\sigma^2_{\mu_t} = \left(\frac{N}{\langle\sigma_t^2\rangle}\right)^{-1}$$

$$\mu_{\mu_t} = \sigma^2_{\mu_t}\sum_n \frac{1}{\langle\sigma_t^2\rangle}(r_{nt}^{\mu_t})$$

$$r_{nt}^{\mu_t} = Y_{nt} - \sum_l X_{nl}\langle W_{lt}\rangle - \sum_i I[n \in i]\langle\alpha_{it}\rangle - G_{nt}\langle F_t\rangle - \ G_{nt}\sum_k \langle U_{nk}\rangle\langle V_{kt}\rangle$$

Individual specific random effects intercept ($\alpha_{it}$)

For each individual $i$ and test $t$

Prior distribution: $p(\alpha_{it}) \sim N(0, \psi_t^2)$

Variational distribution: $q(\alpha_{it}) \sim N(\mu_{\alpha_{it}}, \sigma_{\alpha_{it}}^2)$

Where updates are:

$$\sigma_{\alpha_{it}}^2 = \left(\frac{\sum_n I[n \in i] * 1.0}{\langle \sigma_t^2 \rangle} + \frac{1.0}{\psi_t^2}\right)^{-1}$$

$$\mu_{\alpha_{it}} = \sigma_{\alpha\,it}^2 \sum_n \frac{I[n \in i]}{\langle \sigma_t^2 \rangle} (r_{nt}^{\alpha_{it}})$$

$$r_{nt}^{\alpha_{it}} = Y_{nt} - \langle \mu_t \rangle - \sum_l X_{nl} \langle W_{lt} \rangle - G_{nt} \langle F_t \rangle - G_{nt} \sum_k \langle U_{nk} \rangle \langle V_{kt} \rangle$$

Latent context variance ($\gamma_k^2$)

For each latent context $k$

Prior distribution: $p(1/\gamma_k^2) \sim Gamma(\alpha_0, \beta_0)$

Variational distribution: $q(1/\gamma_k^2) \sim Gamma(\alpha_{\gamma_k^2}, \beta_{\gamma_k^2})$

Where updates are:

$$\alpha_{\gamma_k^2} = \alpha_0 + \frac{N}{2}$$

$$\beta_{\gamma_k^2} = \beta_0 + \frac{\sum_n \langle U_{nk}^2 \rangle}{2}$$

Random effects intercept variance ($\psi_t^2$)

For each test $t$

Prior distribution: $p(1/\psi_t^2) \sim Gamma(\alpha_0, \beta_0)$

Variational distribution: $q\left(\frac{1}{\psi_t^2}\right) \sim Gamma\left(\alpha_{\psi_t^2}, \beta_{\psi_t^2}\right)$

Where updates are:

$$\alpha_{\psi_t^2} = \alpha_0 + \frac{I}{2}$$

$$\beta_{\psi_t^2} = \beta_0 + \frac{\sum_i \langle \alpha_{it}^2 \rangle}{2}$$

<u>Residual variance ($\sigma_t^2$)</u>

For each test $t$

Prior distribution: $p(1/\sigma_t^2) \sim Gamma(\alpha_0, \beta_0)$

Variational distribution: $q\left(\frac{1}{\sigma_t^2}\right) \sim Gamma(\alpha_{\sigma_t^2}, \beta_{\sigma_t^2})$

Where updates are:

$$\alpha_{\sigma_t^2} = \alpha_0 + \frac{N}{2}$$

$$\beta_{\sigma_t^2} = \beta_0 + \frac{1}{2}\sum_n \langle \left( Y_{nt} - \mu_t - \sum_i I[n \in i]\langle \alpha_{it} \rangle - \sum_l X_{nl} W_{lt} - G_{nt}F_t - G_{nt}\sum_k U_{nk} V_{kt} \right)^2 \rangle$$

**SURGE coordinate ascent variational inference update algorithm**

Below, we provide pseudocode documenting the SURGE inference algorithm:

- Randomly initialize variational distributions $q(Z)$, where [$Z = (F_t, V_{kt}, U_{nk}, \mu_t, W_{lt}, \alpha_{it}, \gamma_k^2, \psi_t^2, \sigma_t^2)$]
- iteration_num = 0
- While inference has not converged:
    - Update $q(U_{nk})\ \forall\ n, k$
    - Update $q(V_{kt})\ \forall\ k, t$

- Update $q(\alpha_{it})\ \forall\ i,t$
- Update $q(C_{lt})\ \forall\ l,t$
- Update $q(F_t)\ \forall\ t$
- If iteration_num >=5:
    - Update $\gamma_k^2\ \forall\ k$
- Update $q(\psi_t^2)\ \forall\ t$
- Update $q(\sigma_t^2)\ \forall\ t$
- iteration_num=iteration_num+1
- Converge if change in ELBO < $1e^{-2}$

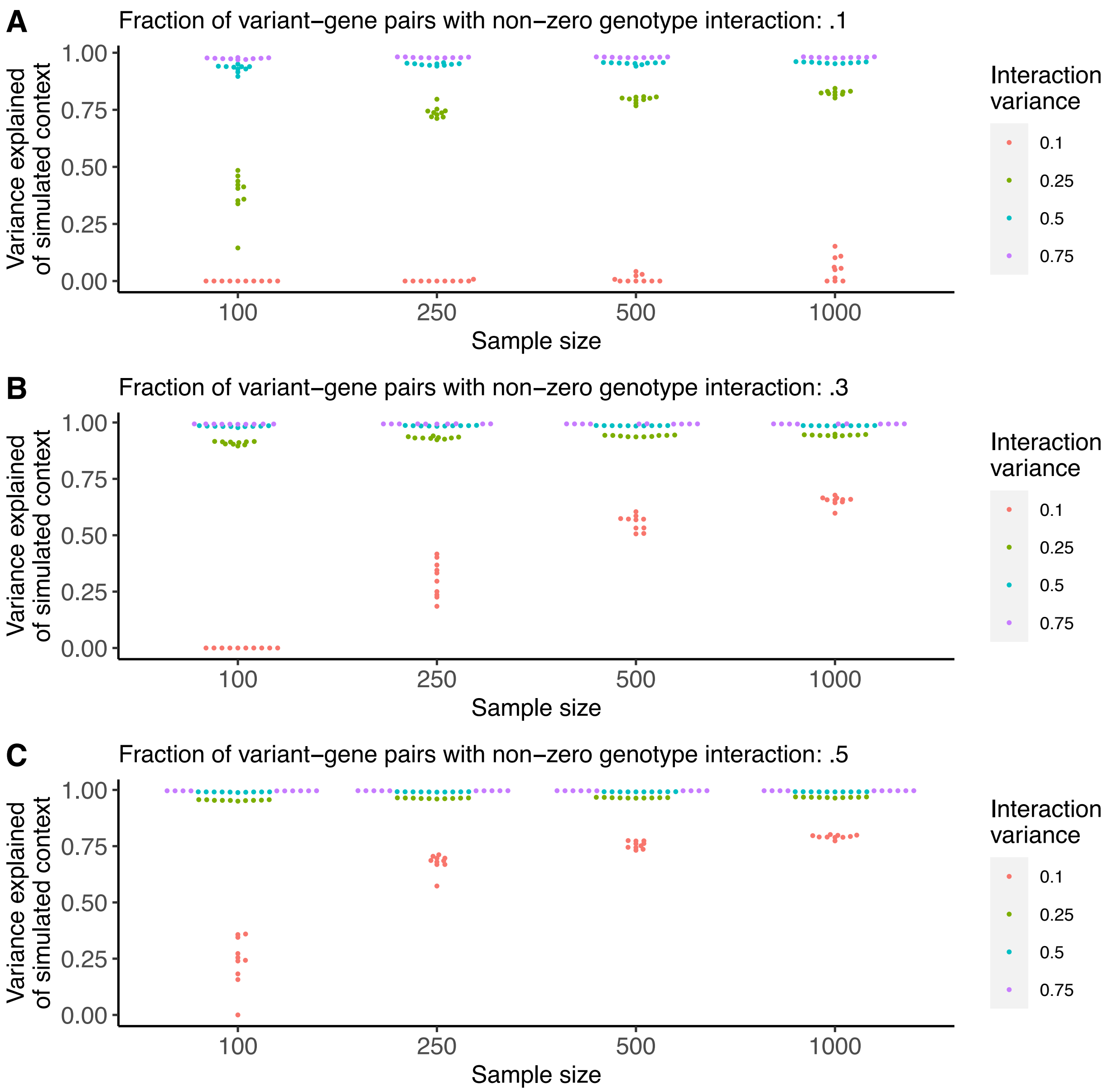


*Figure S1: In this simulation, we evaluate SURGE's ability to re-capture simulated latent contexts as measured by the variance explained of the simulated components by the learned components (y-axis). In this simulation we vary the sample size (x-axis), the strength (variance) of the simulated interaction terms (colors), and the fraction of tests that are context-specific eQTLs for a particular context (A, B, C). For each parameter setting, we run 10 independent simulations. Each dot represents an independent simulation.*

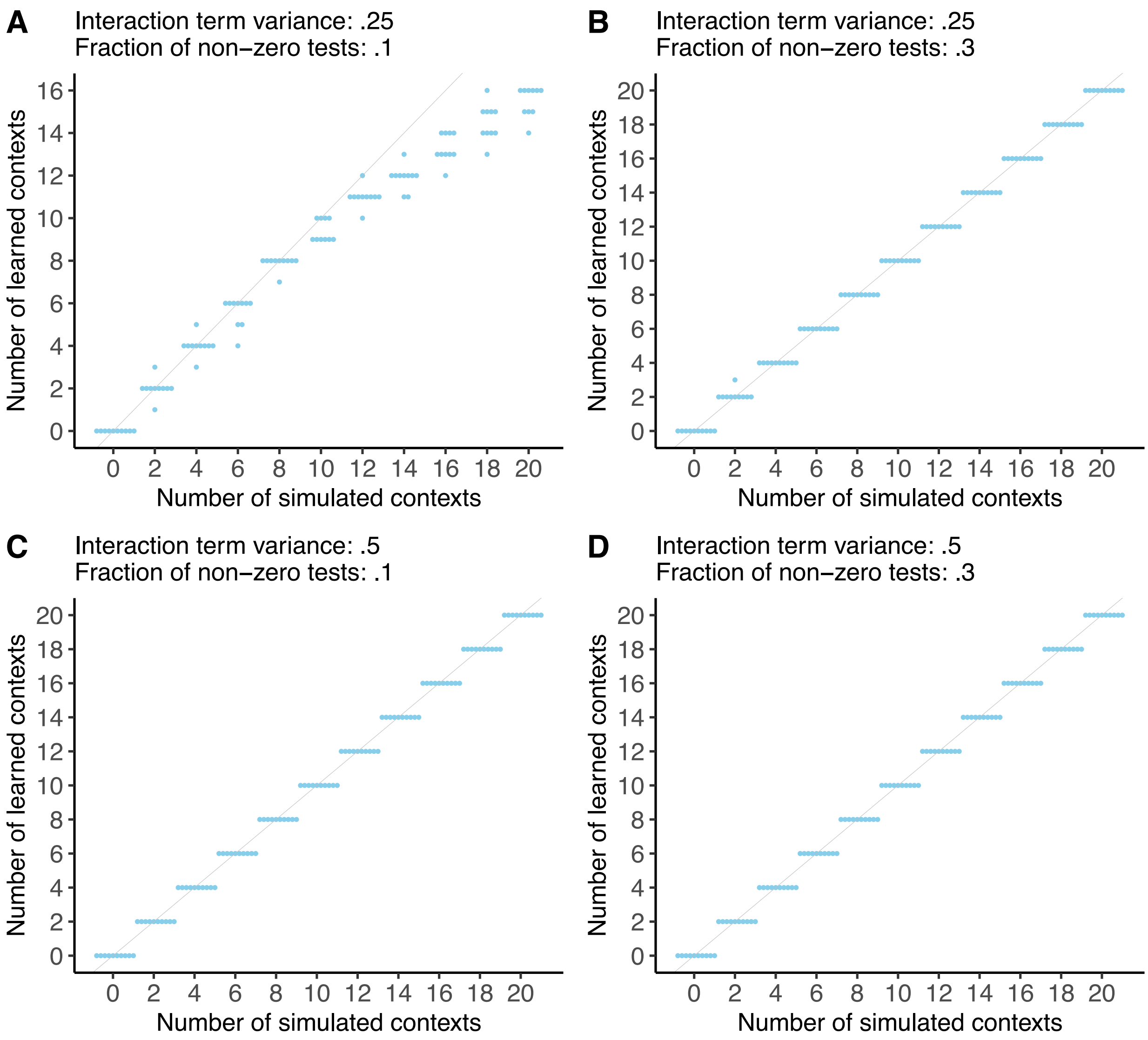


*Figure S2: In this simulation, we evaluate SURGE's ability to identify the number of simulated latent contexts (x-axis) over 10 independent simulations and SURGE optimizations (y-axis). In this simulation, the sample size was fixed to 250, the strength (variance) of the simulated interaction terms was set to .25 (A, B) and .5 (C, D), and the fraction of tests that are context-specific eQTLs for a particular context was set to .1 (A, C) and .3 (B,D).*

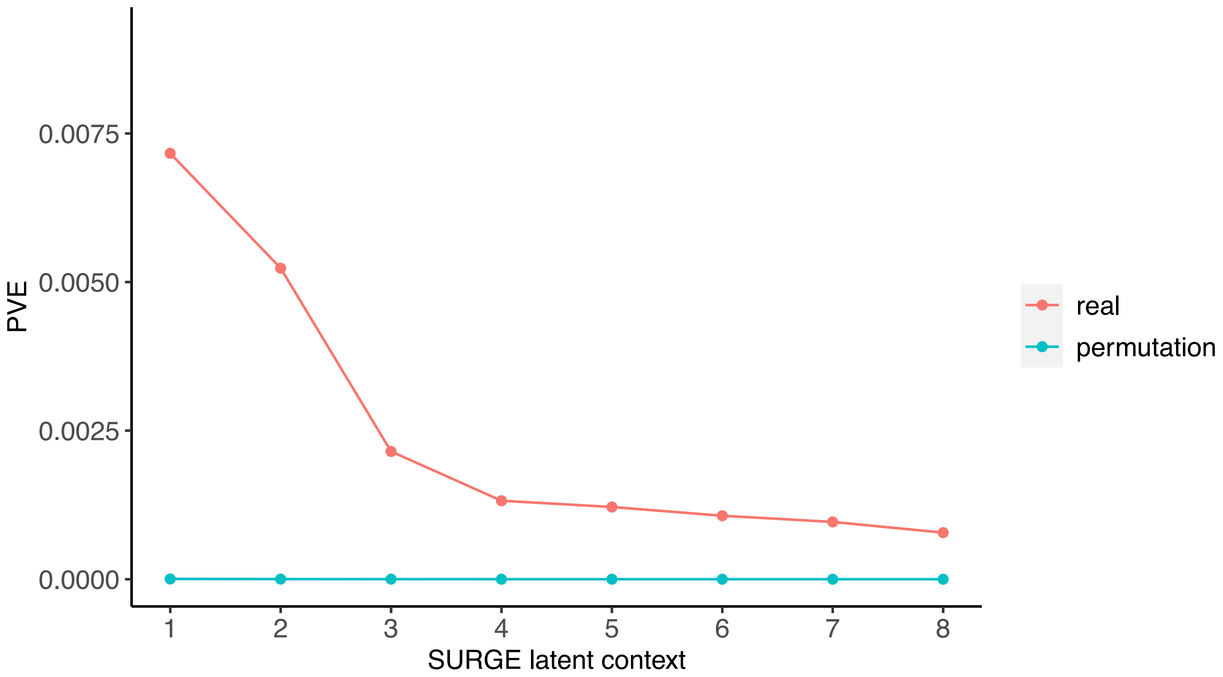


*Figure S3: Percent variance explained (PVE; see Methods; y-axis) of the 8 SURGE latent contexts identified when SURGE was applied to samples concatenated across 10 GTEx v8 tissues. We show results for real data and data with permuted genotype (colors).*

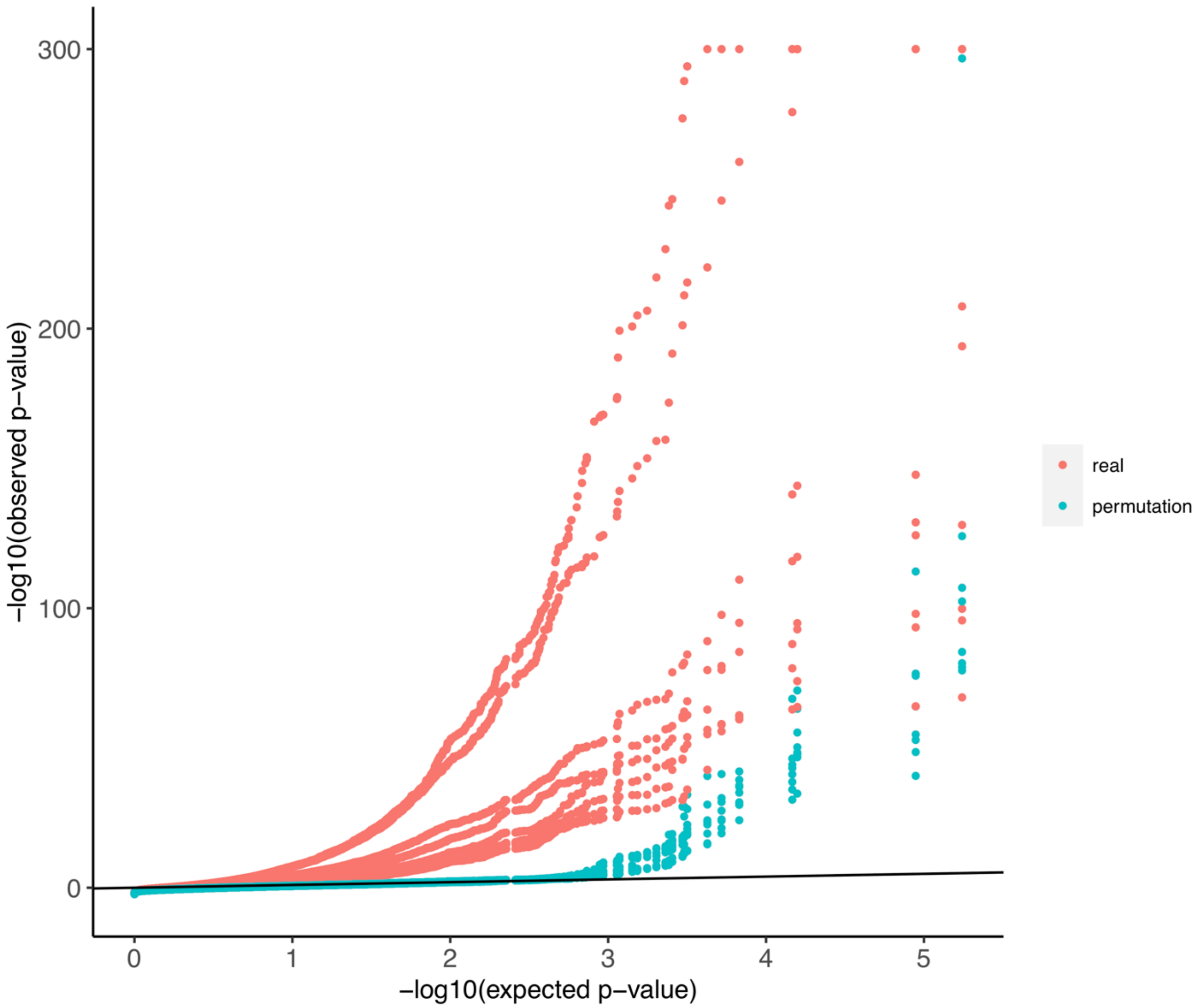


*Figure S4: Q-Q plot for 8 SURGE interaction eQTLs identified in GTEx v8 eQTL data aggregrated across 10 tissues. Red dots correspond to gene-level Bonferonni-corrected p-values for SURGE interaction-eQTLs relative to uniformly distributed p-values. Teal dots correspond to gene-level Bonferonni-corrected p-values from SURGE interaction eQTLs called with permuted genotype relative to uniformly distributed p-values.*

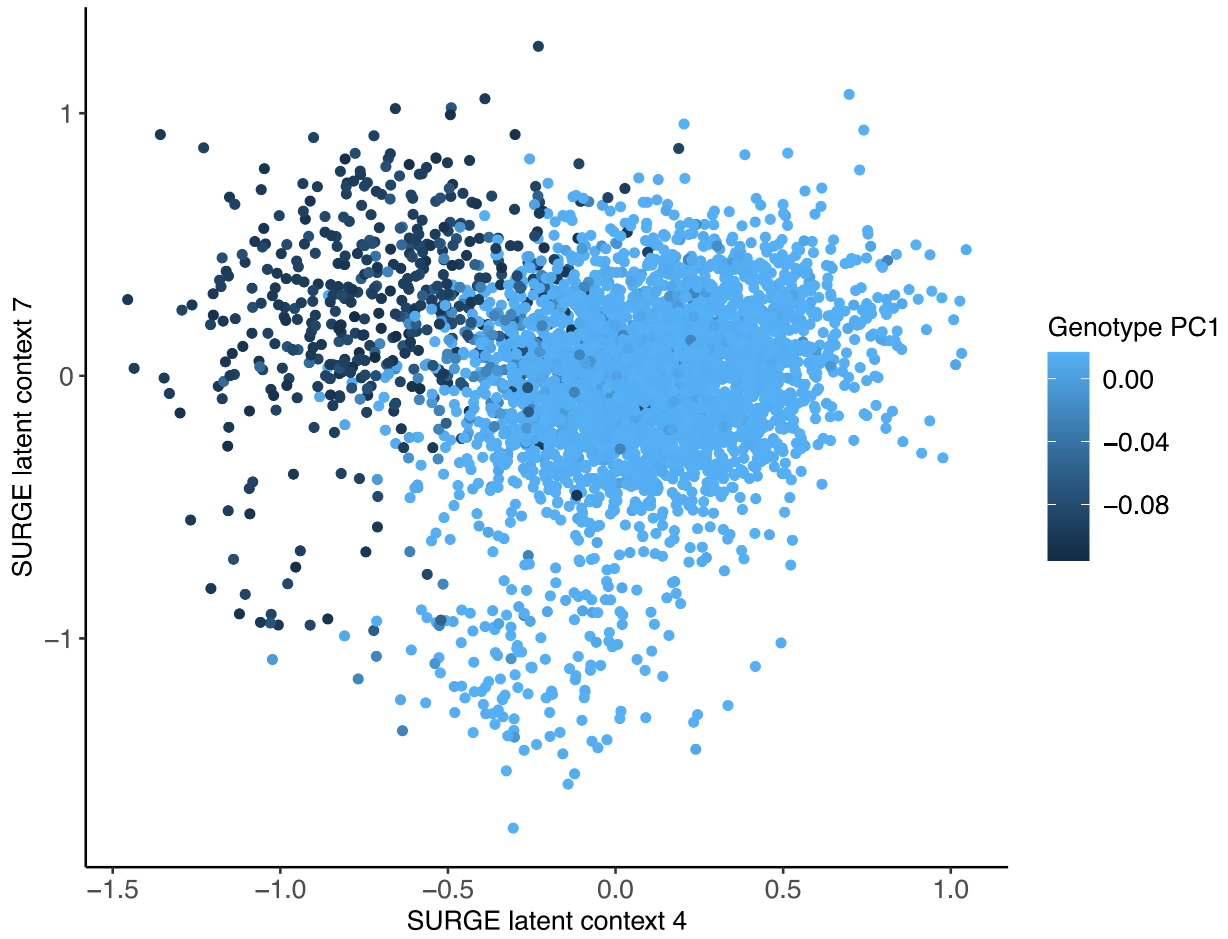


*Figure S5: Scatter-plot of SURGE latent context 4 values (x-axis) by SURGE latent context 7 (y-axis) values across all GTEx version 8 samples concatenated over 10 GTEx tissues. Samples are colored by their loading on the first Genotype PC.*

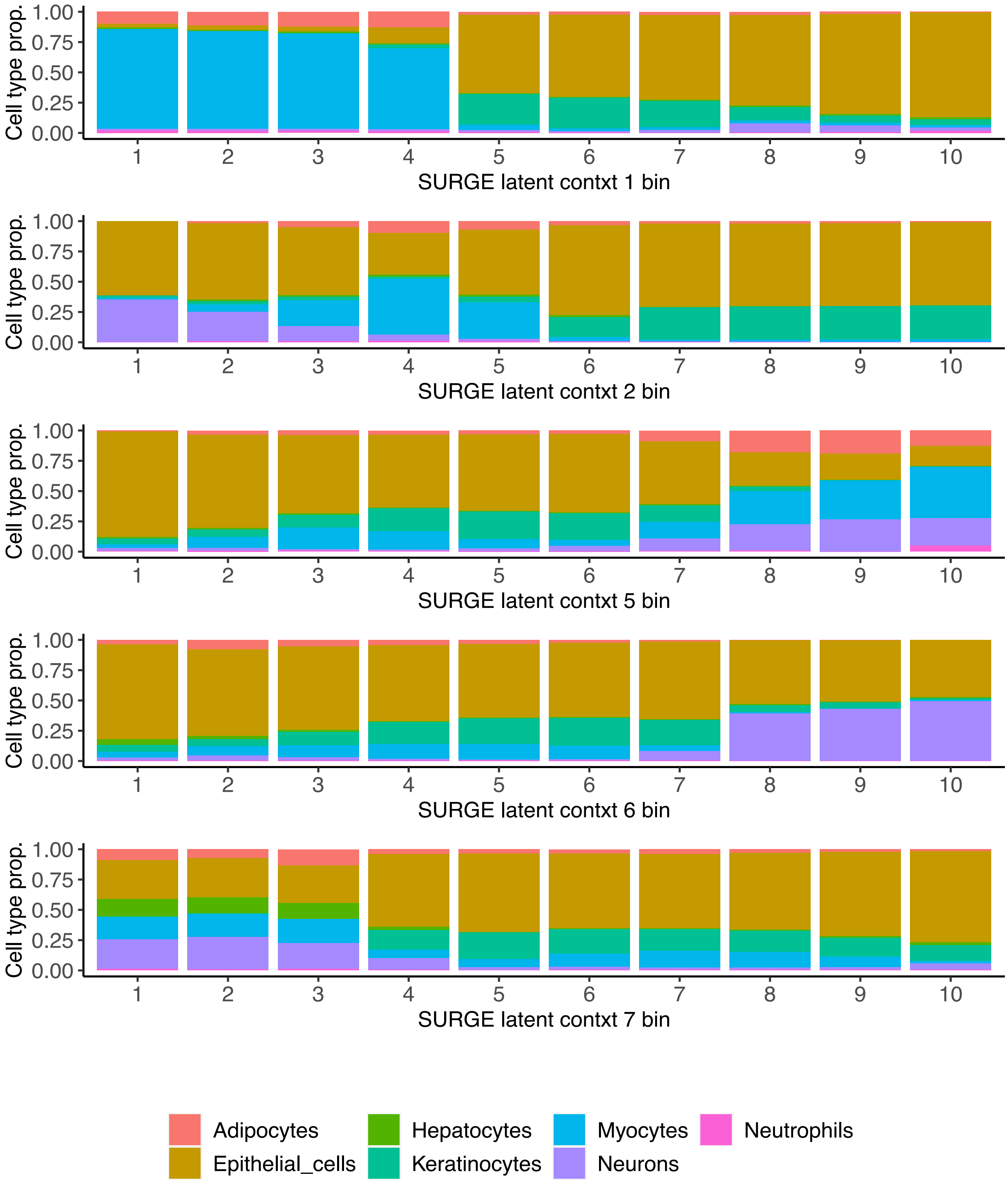


*Figure S6: GTEx v8 RNA-seq samples are separated into 10 equally-sized bins according to their value on SURGE latent context 1, 2, 5, 6, and 7 (rows). The stacked bar plots depicts the average cell-type composition according to xCell estimates across all samples normalized to sum to 1(y-axis) in each of the 10 bins (x-axis). These results were generated when SURGE was applied to samples from 10 GTEx v8 tissues.*

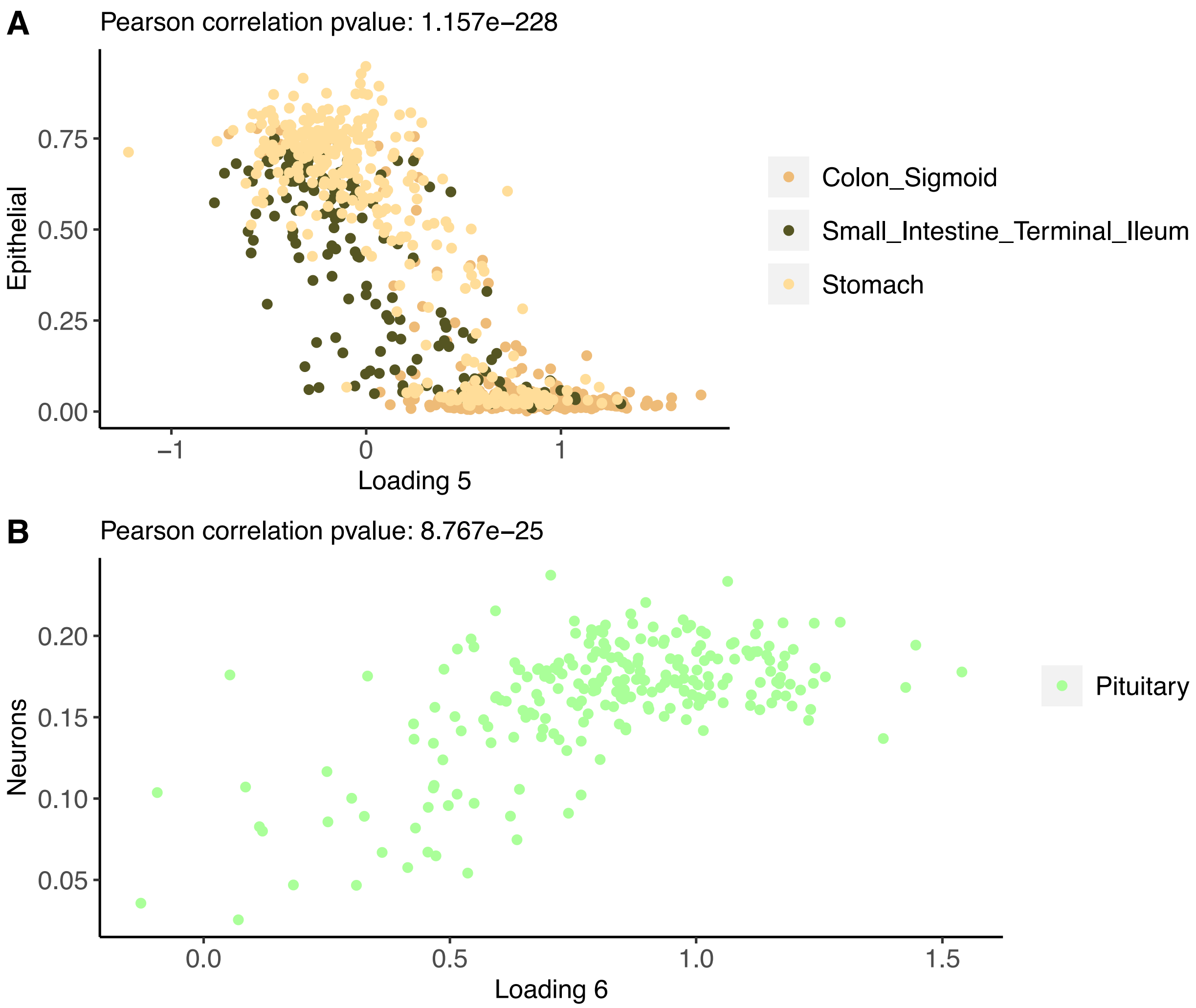


*Figure S7: (A) Scatter-plot of SURGE latent context 5 values (x-axis) by xCell Epithelial cell-type enrichment scores (y-axis) for GTEx v8 samples from Colon Sigmoid, Small Intestine Terminal Ileum, and Stomach tissues. (B) Scatter-plot of SURGE latent context 6 values (x-axis) by xCell Neuron cell-type enrichment scores (y-axis) for GTEx v8 samples from Pituitary tissue.*

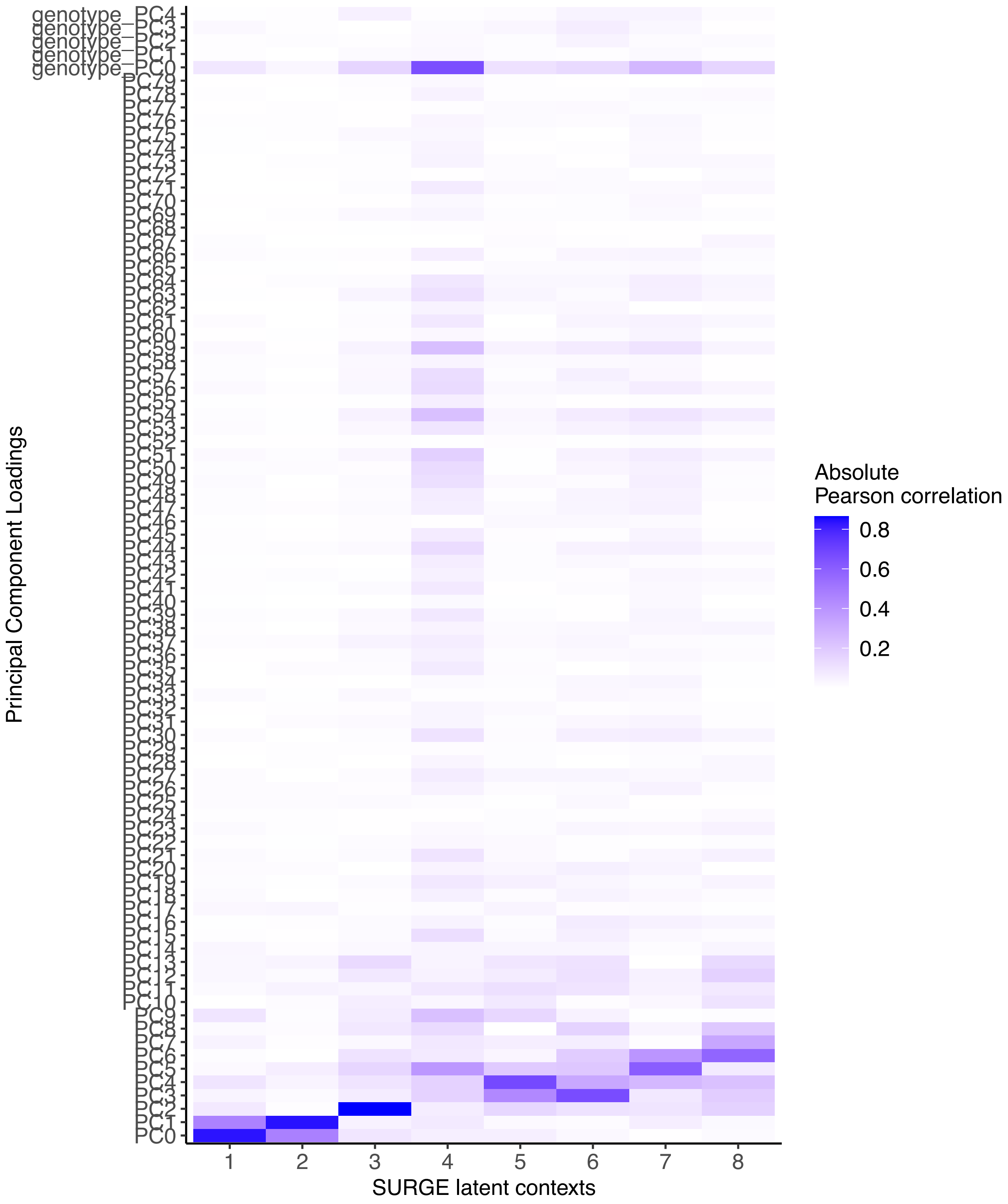


*Figure S8: Heatmap showing absolute Pearson correlation between the 8 SURGE latent contexts (identified when run on 10 GTEx tissues) (x-axis) and gene expression principal components and genotype principal components (y-axis).*

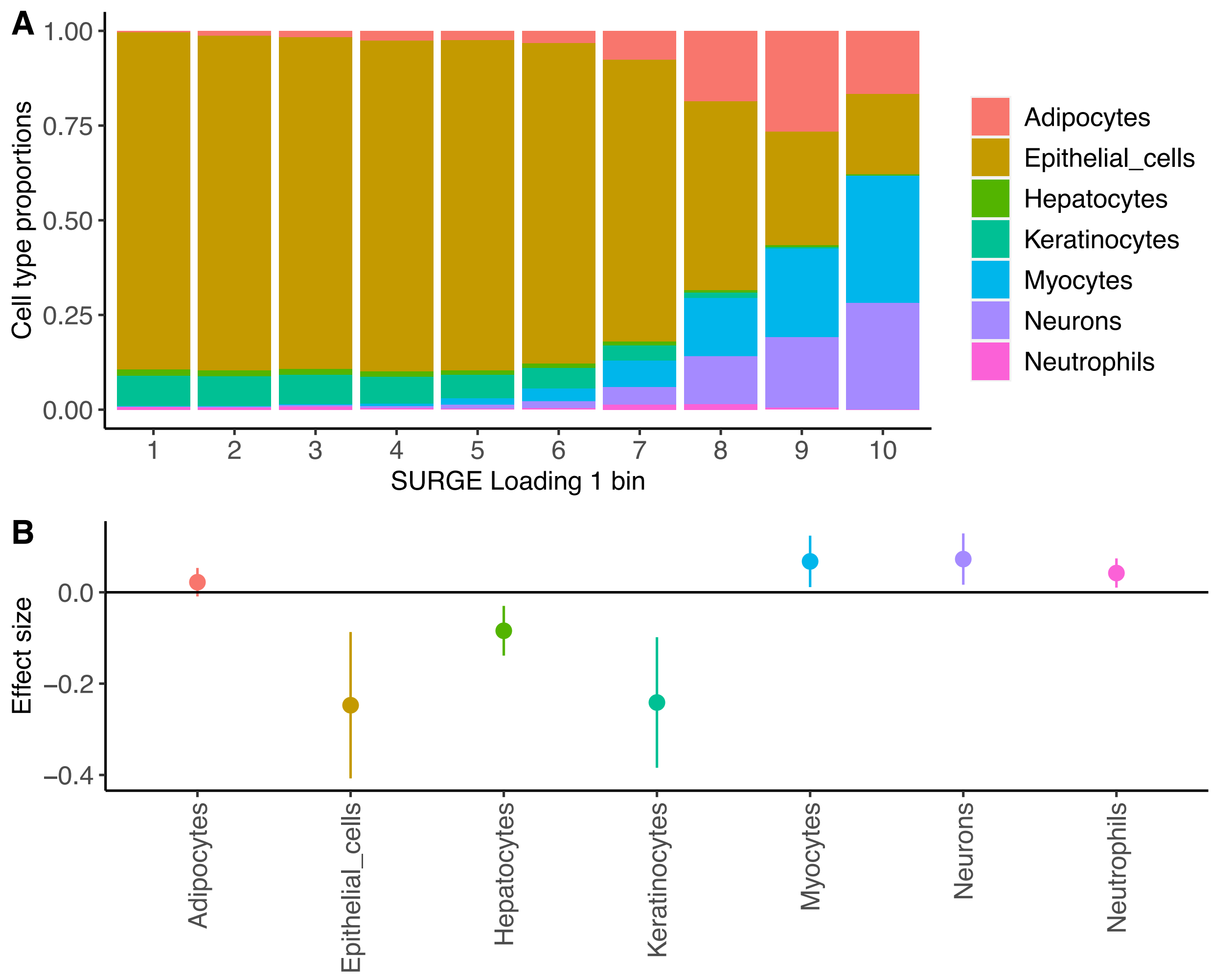


*Figure S9: These results were generated when SURGE was applied to samples from only Colon-Sigmoid GTEx v8 tissue. (A) GTEx v8 Colon-Sigmoid RNA-seq samples are separated into 10 equally-sized bins according to their value on SURGE latent context 1. The stacked bar plot depicts the average cell-type composition according to xCell enrichment scores across all samples normalized to sum to 1 (y-axis) in each of the 10 bins (x-axis). (B) We fit a multivariate linear model to predict SURGE latent context 1 from xCell cell type enrichment scores across 8 cell types. This plot shows the effect sizes and standard error of the effect sizes from this multivariate linear model (y-axis) for each of the 8 cell types that were used as fixed effects in the model (x-axis). 6 of the 7 cell types are predictive of SURGE latent context 1, even when conditioned on all other cell types.*

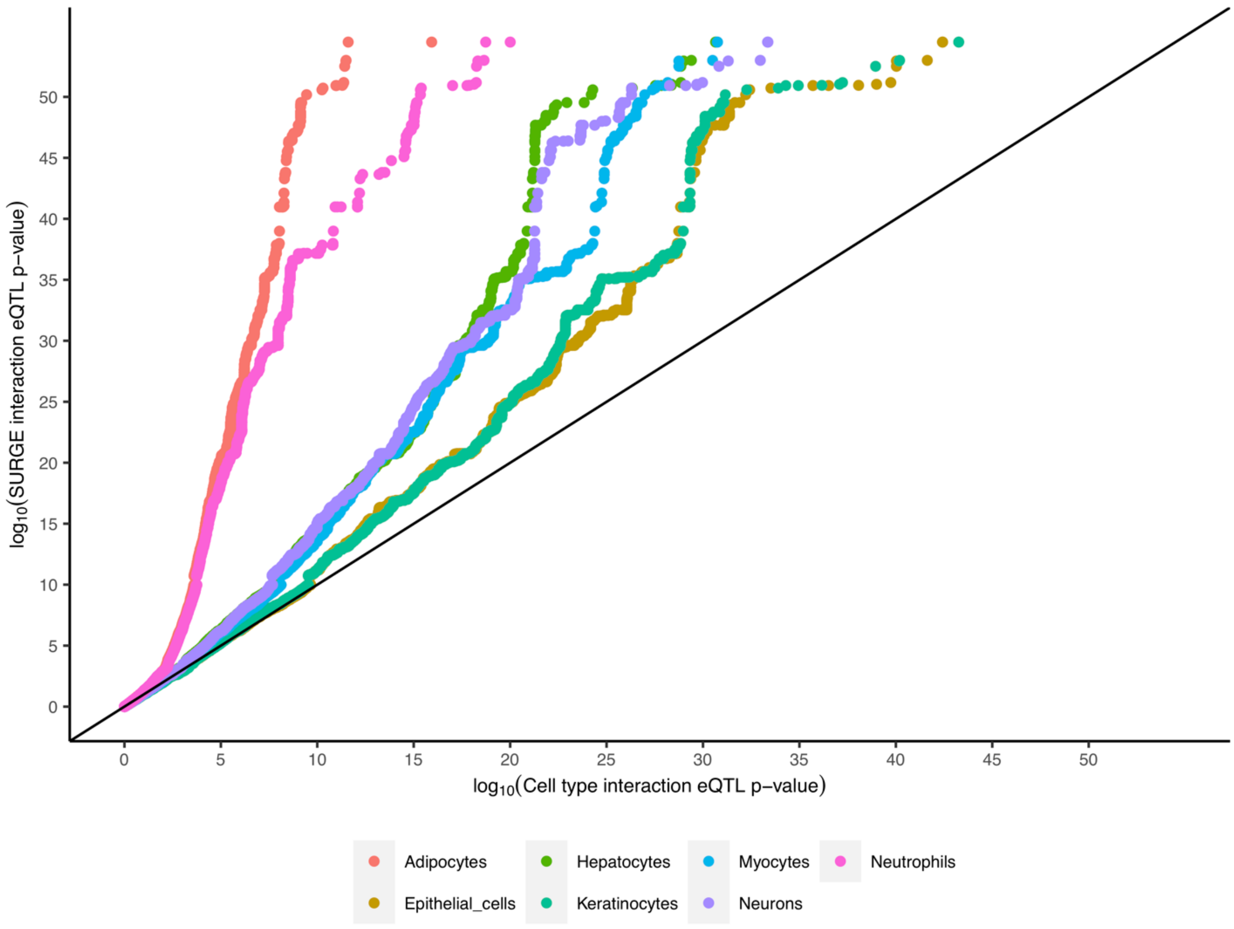


*Figure S10: These results were generated when SURGE was applied to samples from only Colon-Sigmoid GTEx v8 tissue. -log10(pvalues) of SURGE context 1 interaction eQTLs (y-axis) compared to -log10(pvalues) of interaction eQTLs using xCell cell type proportion from single cell type as the context (x-axis). Results shown for all 7 xCell cell types (colors).*

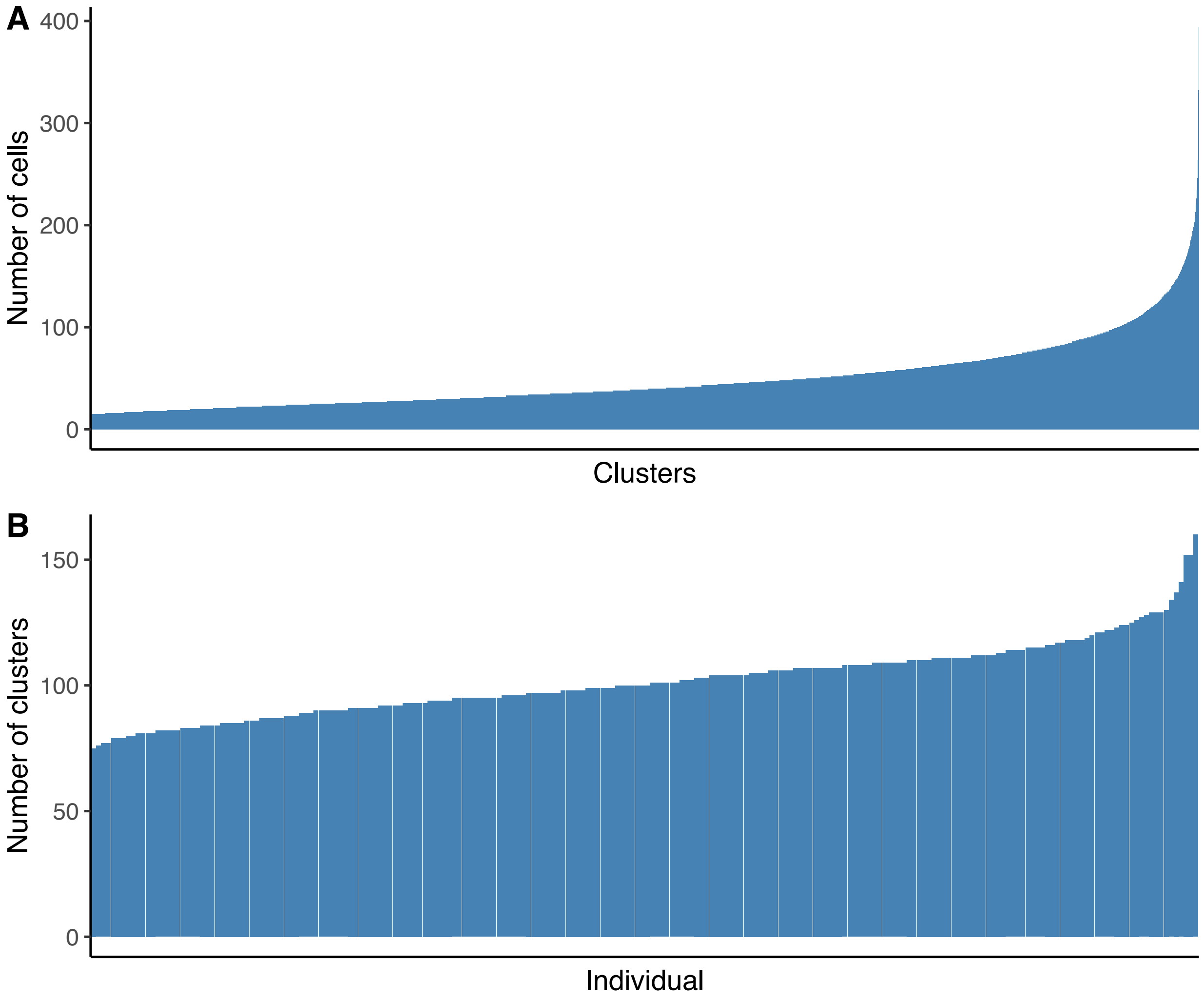


*Figure S11: Pseudocell aggregation of PBMC single cell expression data. (A). Distribution of number of cells (y-axis) per pseuodcell (x-axis). (B) Distribution of number pseudocells (y-axis) per individual (x-axis).*

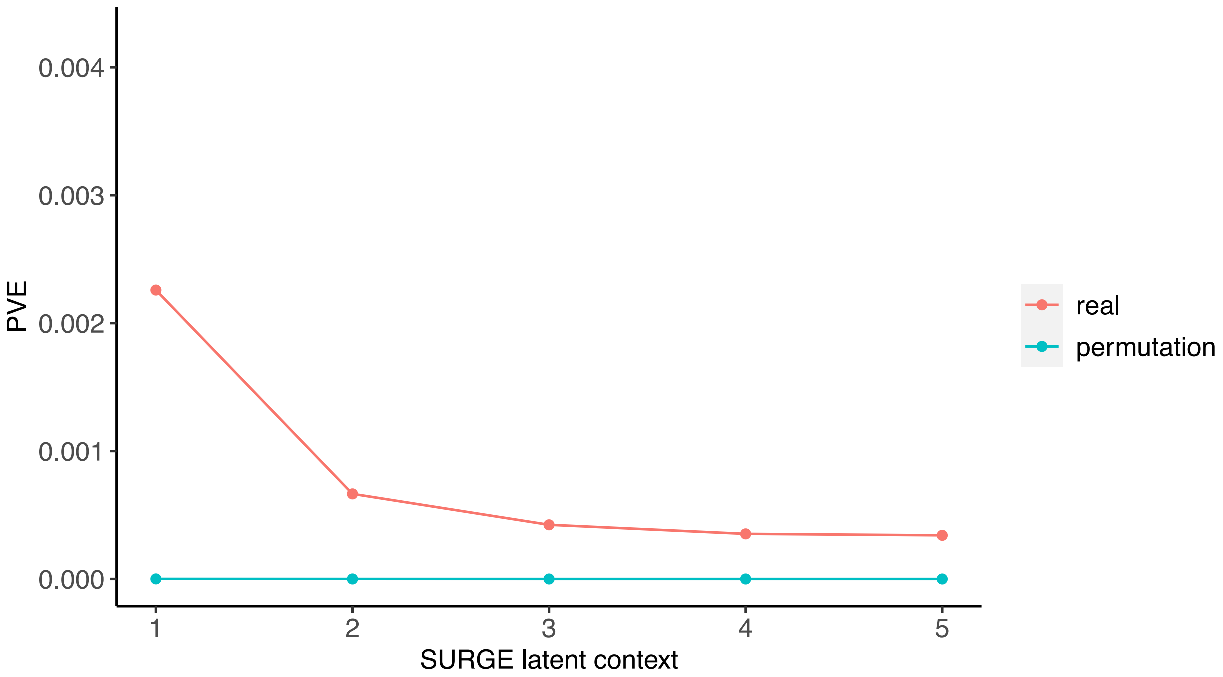


*Figure S12: Percent variance explained (PVE; see Methods; y-axis) of the 5 SURGE latent contexts identified when SURGE was applied to PBMC pseudocells. We show results for real data and data with permuted genotype (colors).*

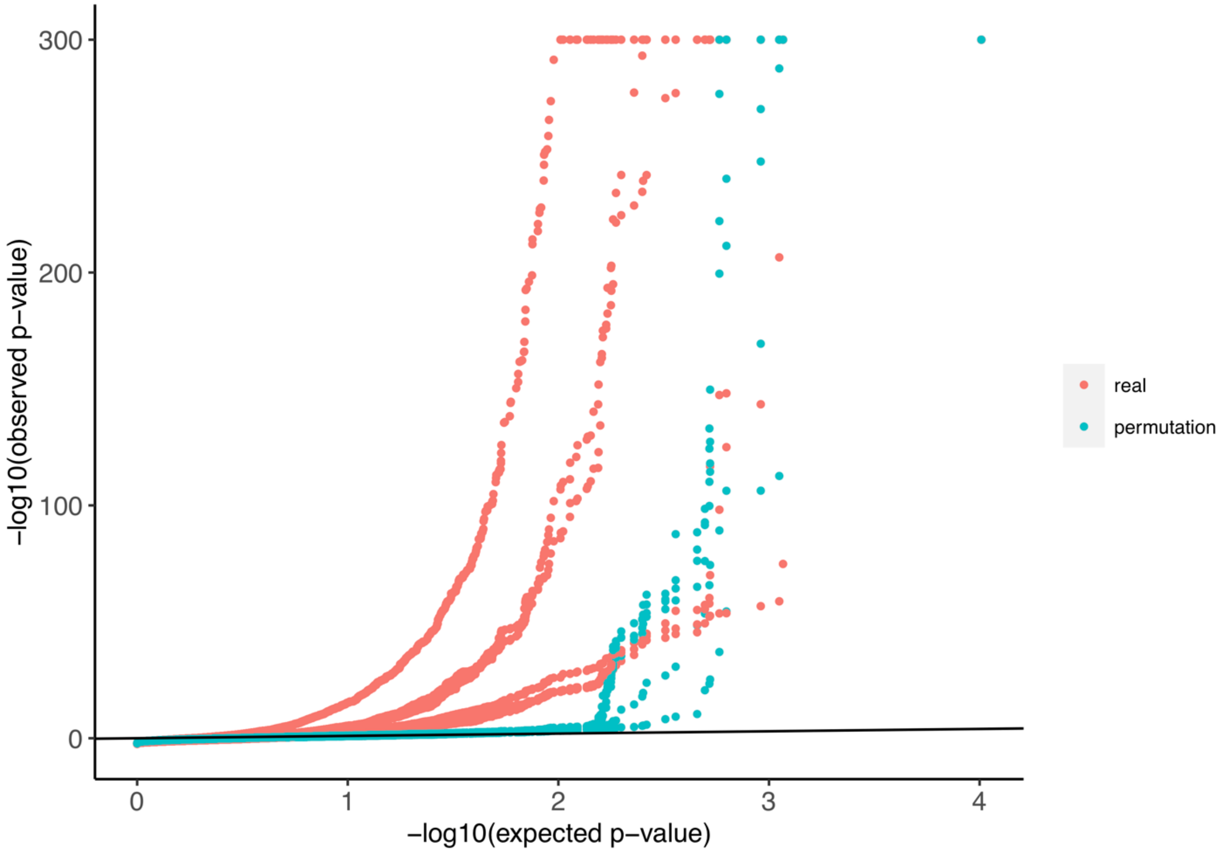


*Figure S13: Q-Q plot for 5 SURGE interaction eQTLs identified in PBMC single-cell eQTL data. Red dots correspond to gene-level Bonferonni-corrected p-values for SURGE interaction-eQTLs relative to uniformly distributed p-values. Teal dots correspond to gene-level Bonferonni-corrected p-values from SURGE interaction eQTLs called with permuted genotype relative to uniformly distributed p-values.*

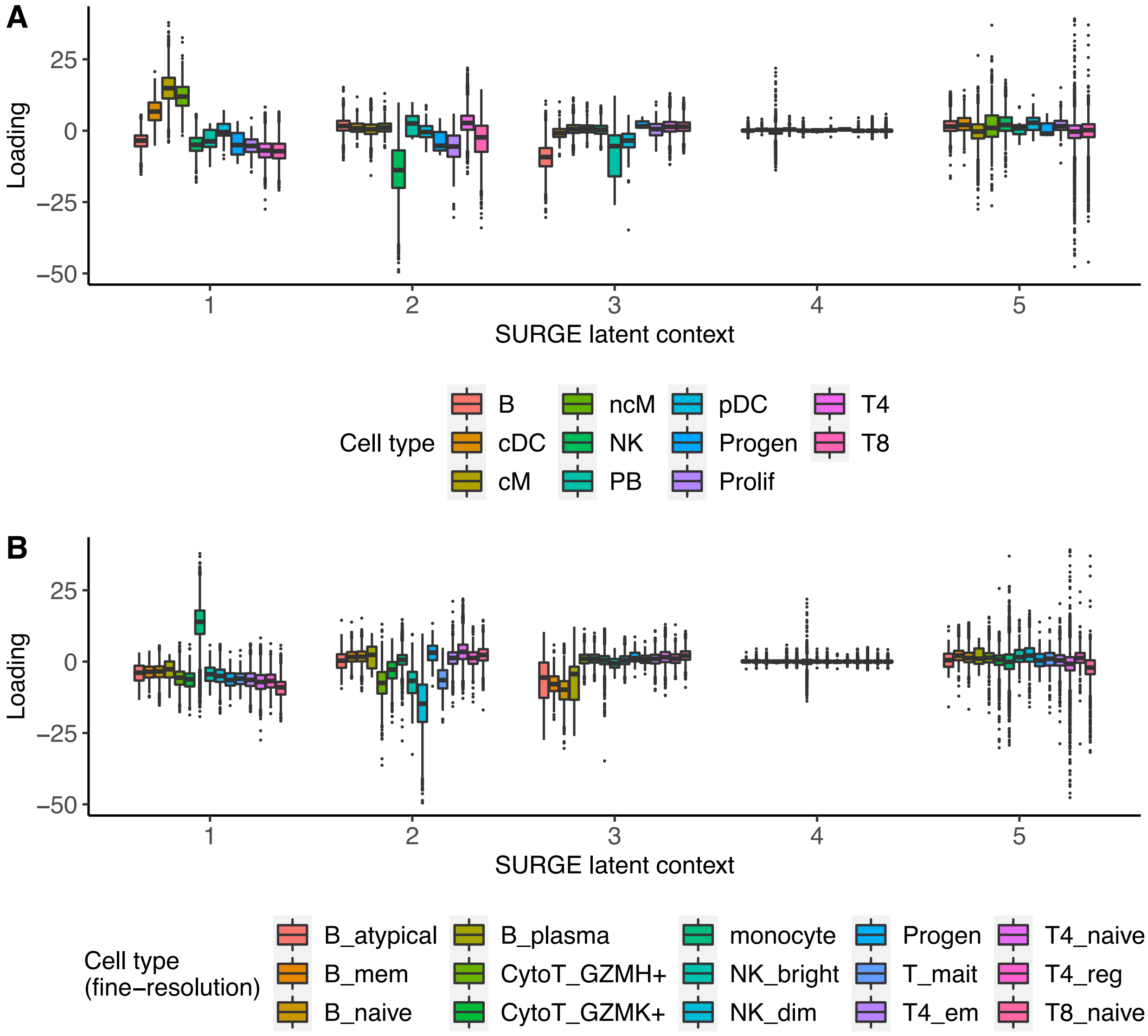


*Figure S14: SURGE latent context loadings of pseudocells (y-axis) stratified by cell type according to marker gene expression profiles for each of the 5 identified SURGE latent contexts. Results shown for both (A) cell-types and (B) fine-resolution cell-types.*

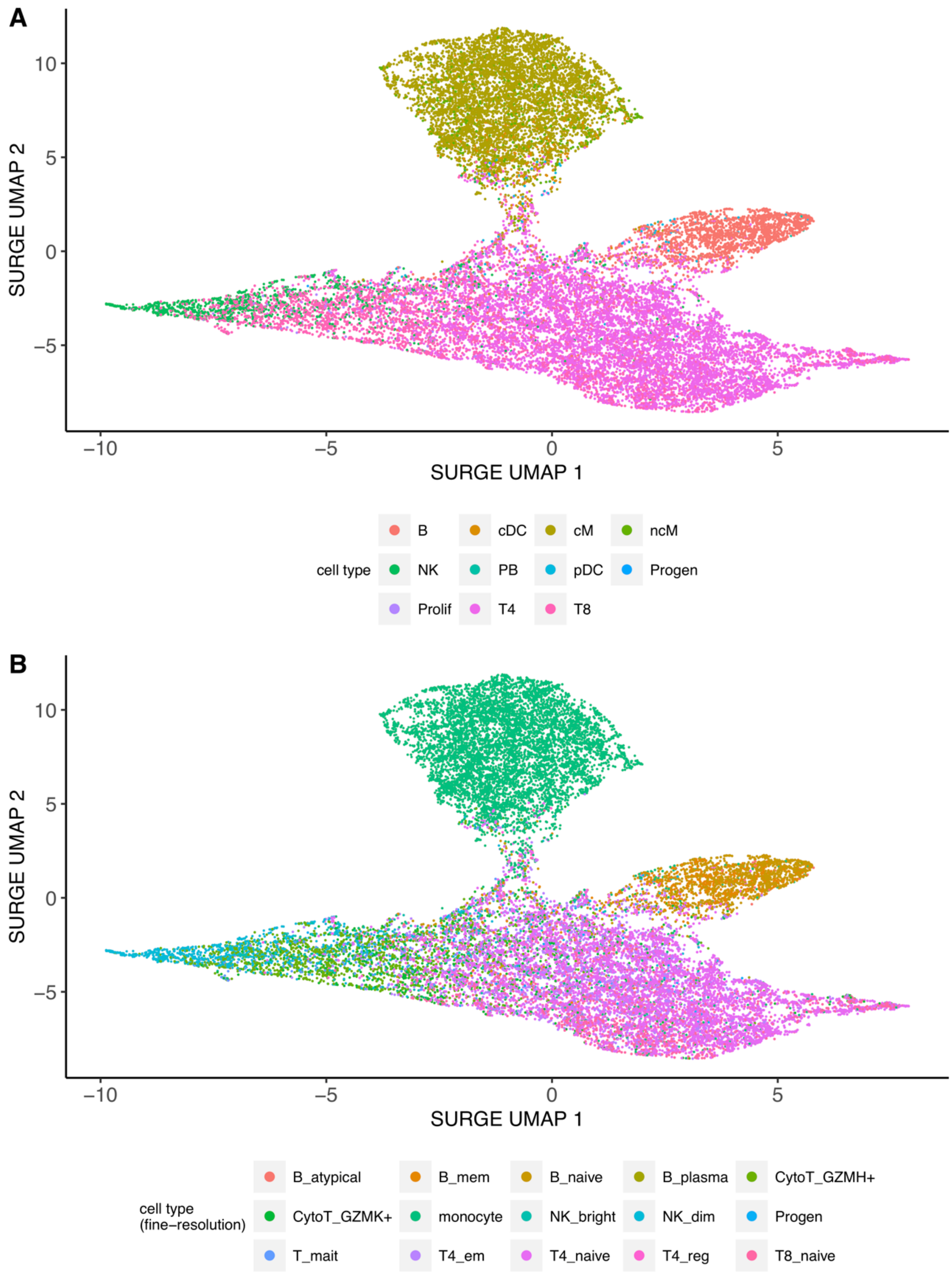


*Figure S15: UMAP-projected SURGE latent context loadings of pseudocells (x and y-axis) colored by (A) cell-types and (B) fine-resolution cell-types.*

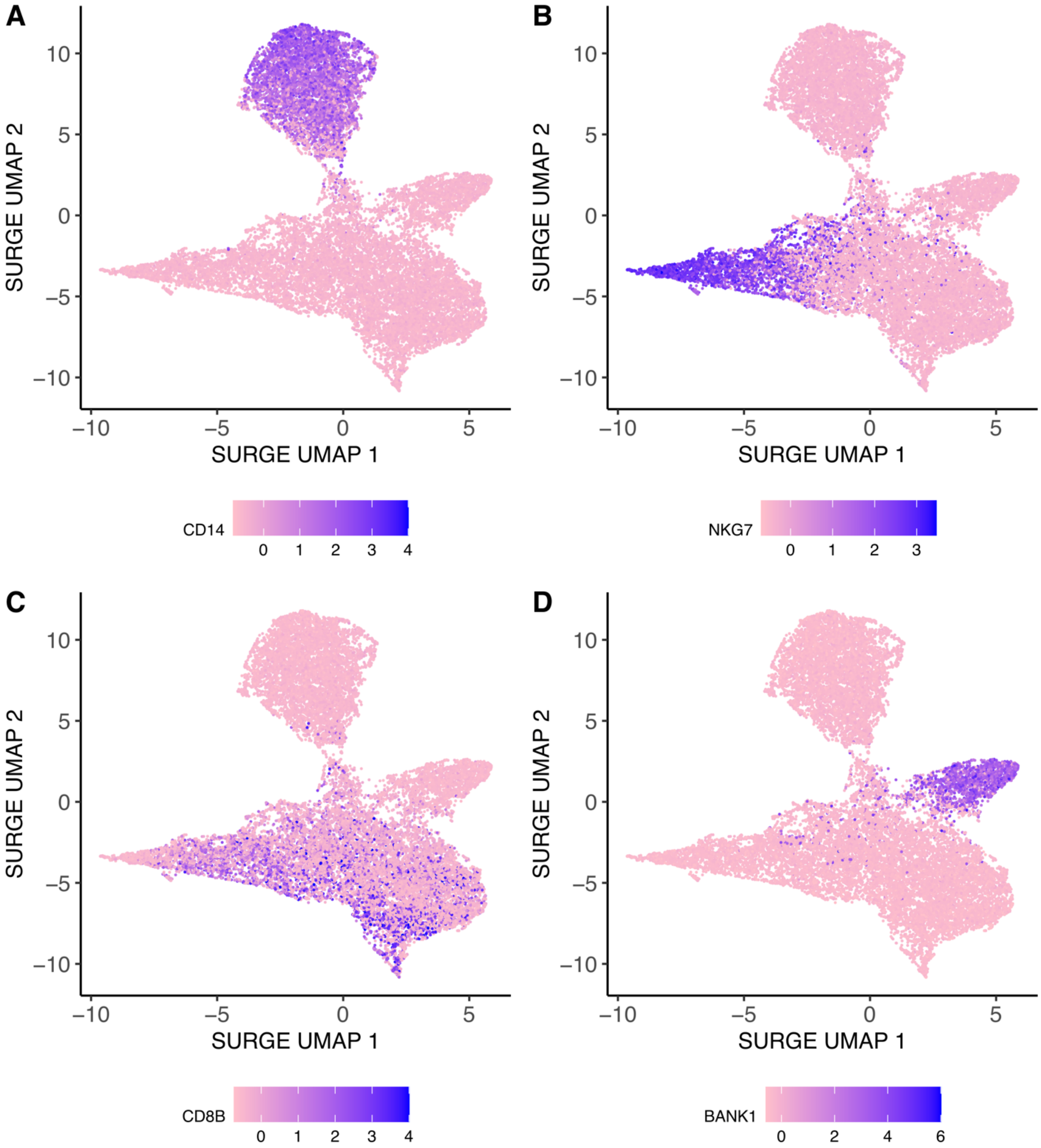


*Figure S16: UMAP-projected SURGE latent context loadings of pseudocells (x and y-axis) colored by expression levels of four marker genes: (A) CD14, (B) NKG7, (C) CD8B, (D) BANK1. CD14 is a marker for monocytes, NKG7 is a maker for NK cells, CD8B is a marker for T cells and BANK1 is a marker for B cells.*

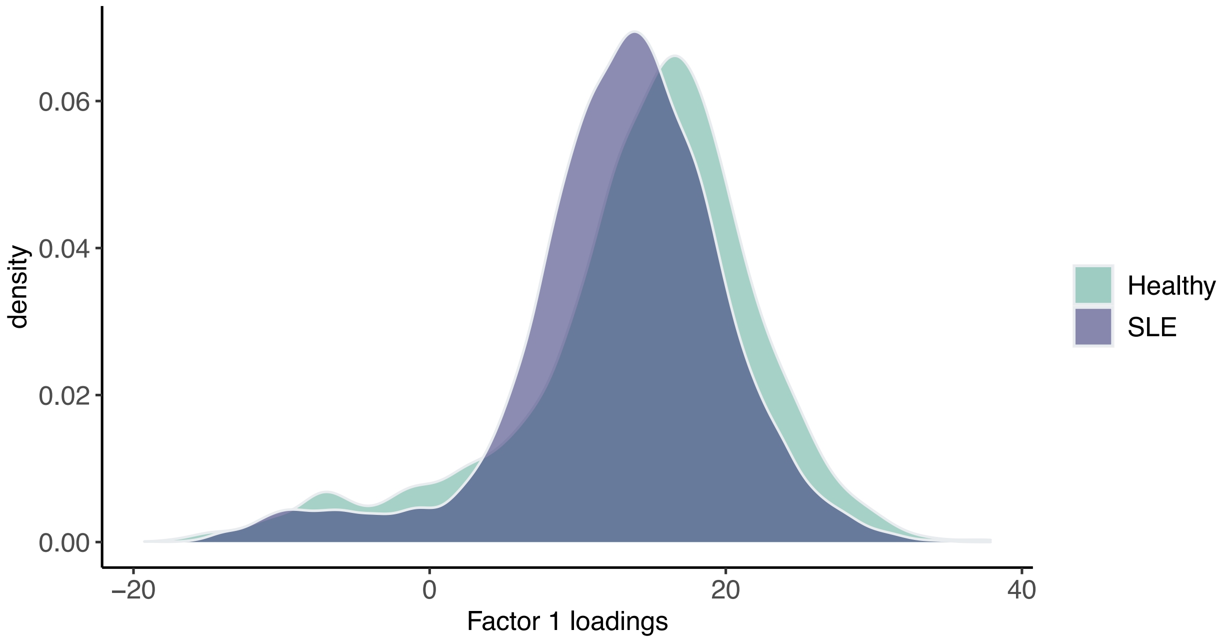


*Figure S17: Density of (y-axis) SURGE latent context 1 loadings (x-axis) on pseudocells annotated as monocytes according to marker-gene expression profiles color-stratified by disease status of individuals corresponding to pseudocells.*

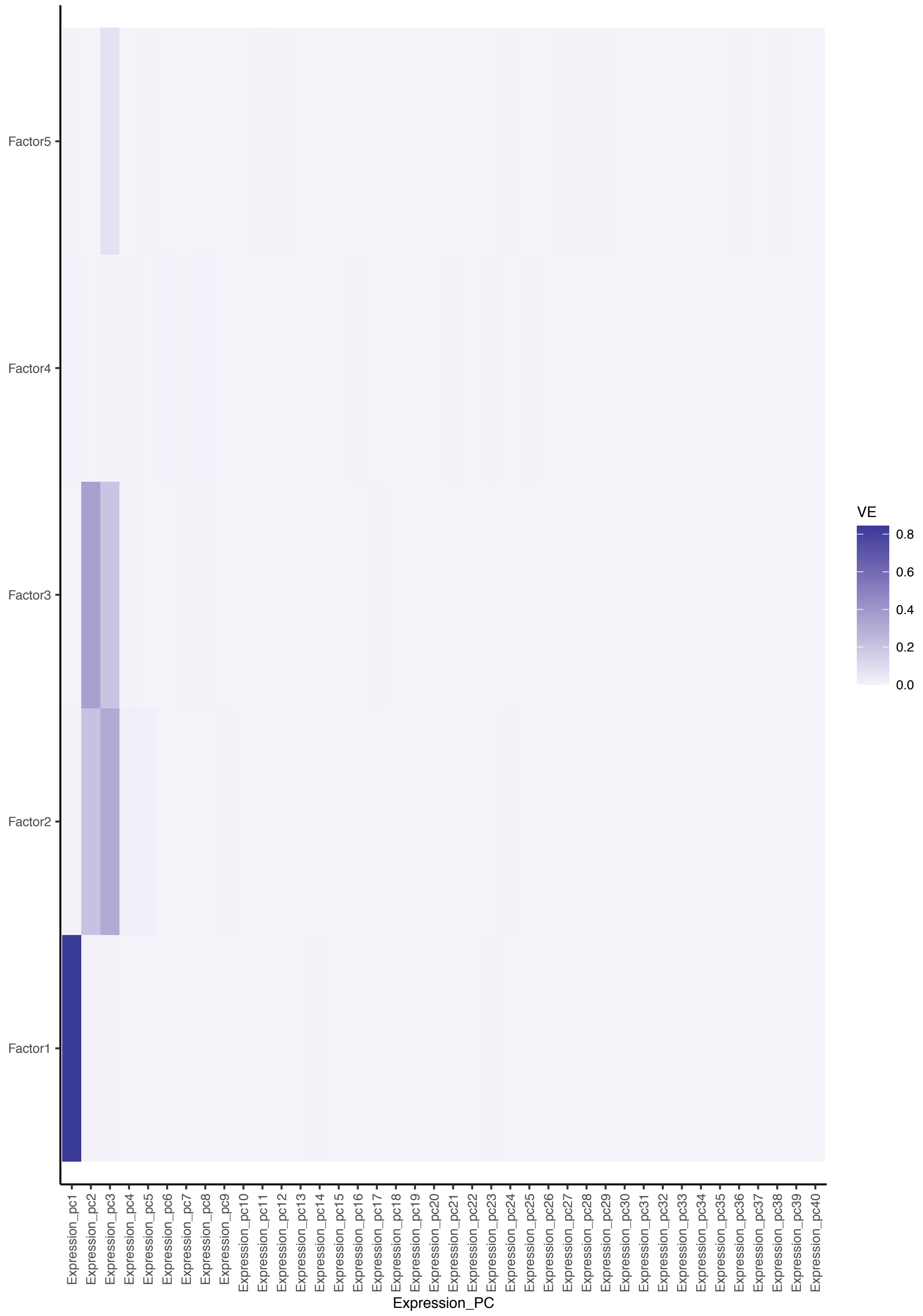


*Figure S18: Heatmap showing variance explained of each of the gene expression principal components (x-axis) by the 5 SURGE latent contexts (y-axis) (identified when run on PBMC single-cell eQTL data set).*

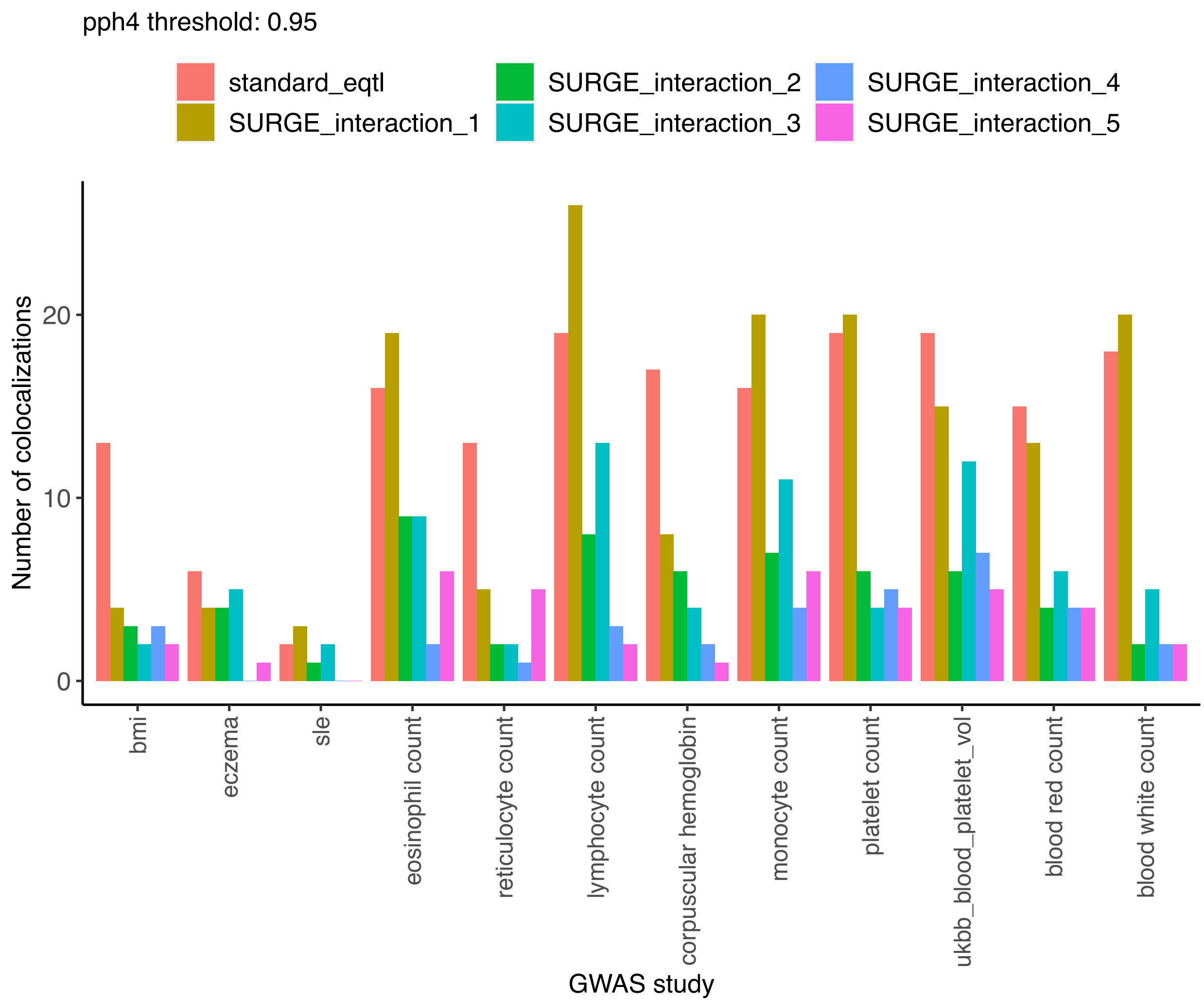


*Figure S19: Number of colocalizations identified (PPH4 > .95; y-axis) between various densely genotyped GWAS studies (x-axis) and various categories of eQTLs called from pseudocells.*

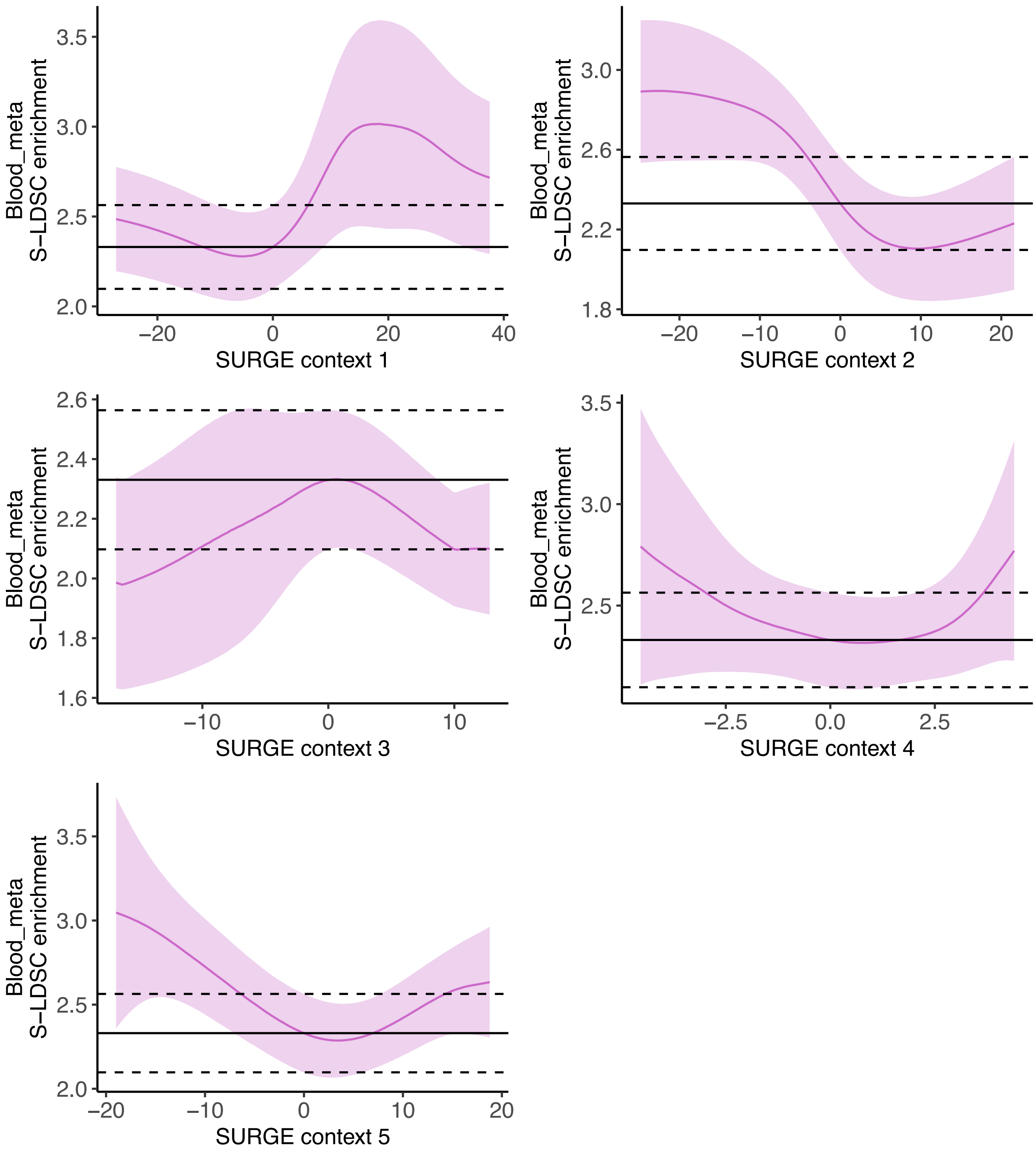


*Figure S*20: S-LDSC enrichment (y-axis) of squared standard eQTL effect sizes (black line) and SURGE predicted squared eQTL effect size at a specific SURGE latent context value (pink line at a specific x-axis position) meta-analyzed across blood-related traits shown for 5 SURGE latent contexts. SURGE predicted eQTL effect sizes at a particular SURGE latent context value was calculated at 200 equally spaced positions along the range of SURGE latent context values. Black dashed line represents 95% confidence on the standard eQTL S-LDSC enrichment. Light pink region depects 95% confidence on the SURGE predicted eQTL S-LDSC enrichment. Trait category of “blood” consists of GWAS for eosinophil count, reticulocyte count, lymphocyte count, corpuscular hemoglobin, monocyte count, platelet count, blood platelet volume, red blood count, and white blood count.

| SURGE latent context | Number of genes with SURGE interaction-eQTL (eFDR < .05) | Number of genes with with SURGE interaction-eQTL (eFDR < .1) |
|---|---|---|
| 1 | 5649 | 6771 |
| 2 | 5336 | 6335 |
| 3 | 2363 | 3223 |
| 4 | 4261 | 5677 |
| 5 | 1311 | 1872 |
| 6 | 985 | 1672 |
| 7 | 1087 | 1682 |
| 8 | 587 | 1187 |

*Table S1: The number of genes with a genome-wide significant variant that is a SURGE interaction-eQTL for each of the 8 SURGE latent contexts (rows) identified when SURGE was run on 10 GTEx tissues. Significance determined via empirical FDR (eFDR) correction according to an empirical null distribution generated from a permutation analysis (see Methods).*

| SURGE latent context | Number of genes with SURGE interaction-eQTL (eFDR < .05) | Number of genes with with SURGE interaction-eQTL (eFDR < .1) |
|---|---|---|
| 1 | 1407 | 1641 |
| 2 | 0 | 657 |
| 3 | 514 | 824 |
| 4 | 0 | 297 |
| 5 | 0 | 406 |

*Table S2: The number of genes with a genome-wide significant variant that is a SURGE interaction-eQTL for each of the 5 SURGE latent contexts (rows) identified when SURGE was run on PBMC single cell eQTL data. Significance determined via empirical FDR (eFDR) correction according to an empirical null distribution generated from a permutation analysis (see Methods).*

| ***Hallmark gene set*** | ***Latent context 4*** | ***Latent context 5*** |
|---|---|---|
| *Interferon gamma response* | *4.40e-12* | *NS* |
| *Interferon alpha response* | *4.19e-5* | *NS* |
| *IL2 Stat5 signaling* | *.00409* | *NS* |
| *Complement* | *.00600* | *NS* |
| *Hypoxia* | *.00654* | *NS* |
| *Coagulation* | *.0408* | *NS* |
| *Allograft rejection* | *NS* | *0.00247* |

*Table S3: Bonferroni corrected p-values (Fisher's exact) from gene set enrichment of genes strongly correlated with Latent Context 4 and 5 (columns) within Hallmark gene sets (rows). Only gene sets with significant enrichment (Bonferroni p-value <= .05) in genes strongly correlated with latent context 4 or latent context 5 are shown. NS means not significant (Bonferroni p-value > .05).*

| ***MSigDB Biological Process gene set*** | ***Latent context 4*** | ***Latent context 5*** |
|---|---|---|
| *Translation* | *NS* | *3.48e-7* |
| *Cellular biosynthetic process* | *NS* | *2.44e-5* |
| *Biosynthetic process* | *NS* | *8.29e-5* |
| *Cell structure disassembly during apoptosis* | *NS* | *.0308* |
| *Immune response* | *NS* | *.0377* |
| *Immune system process* | *NS* | *.0430* |

*Table S4: Bonferroni corrected p-values (Fisher's exact) from gene set enrichment of genes strongly correlated with Latent Context 4 and 5 (columns) within MSigDB Biological process gene sets (rows; c5.bp.v5.1). Only gene sets with significant enrichment (Bonferroni p-value <= .05) in genes strongly correlated with latent context 4 or latent context 5 are shown. NS means not significant (Bonferroni p-value > .05).*
